# Molecular and Anatomical Characterization of Parabrachial Neurons and Their Axonal Projections

**DOI:** 10.1101/2022.07.13.499944

**Authors:** Jordan L. Pauli, Jane Y. Chen, Marcus L. Basiri, Sekun Park, Matthew E. Carter, Elisendra Sanz, G. Stanley McKnight, Garret D. Stuber, Richard D. Palmiter

## Abstract

The parabrachial nucleus (PBN) is a major hub that receives sensory information from both internal and external environments. Specific populations of PBN neurons are involved in behaviors including food and water intake, pain sensation, breathing regulation, as well as learning and responding appropriately to threatening stimuli. However, it is unclear how many PBN neuron populations exist and how different behaviors may be encoded by unique signaling molecules or receptors. Here we provide a repository of data on the molecular identity, spatial location, and projection patterns on dozens of PBN neuron subclusters. Using single-cell RNA sequencing, we identified 21 subclusters of neurons in the PBN and neighboring regions. Multiplexed *in situ* hybridization showed many of these subclusters are localized to distinct PBN subregions. We also describe two major ascending pathways that innervate distinct brain regions by analyzing axonal projections in 21 Cre-driver lines of mice. These results are all publicly available for download and provide a foundation for further interrogation of PBN functions and connections.

## Introduction

The parabrachial nucleus (PBN), located at the junction of the midbrain and pons, relays sensory information from the periphery primarily to the forebrain, thereby playing a major role in informing the brain of both the internal state (interoception) and external conditions (exteroception) to facilitate responses to adverse conditions and help maintain homeostasis.

The inputs and outputs of the PBN have been studied extensively using anterograde and retrograde methods (Fulwiler and Saper, 1984, Gauriau and Bernard, 2002, Krout and Loewy, 2000, Moga et al., 1990, Norgren and Leonard, 1971, Tokita et al., 2009). More recently, these studies have been supplemented using genetically engineered mice and stereotaxic delivery of viruses encoding fluorescent proteins to analyze the afferents and efferent projections of selected subsets of PBN neurons. The vagus transmits signals from internal organs including the gastrointestinal system to the caudal nucleus tractus solitarius (NTS), which then projects to the PBN; thus, detection of visceral signals related to food and malaise depend on this circuit. Other internal organs, muscle, and bone transmit nociceptive signals via intermediary ganglia to the spinal cord and then directly to the PBN. Ascending fibers from the spinal cord relay peripheral temperature and pain signals directly to the PBN, while trigeminal neurons relay these signals from the face. In addition, blood-borne threats to homeostasis are detected by the area postrema and transmitted to the PBN (Zhang et al., 2021). Taste is transmitted from the tongue and palate via branches of the facial, petrosal, glossopharyngeal nerves to the rostral NTS and from there to the PBN. Additionally, calcium imaging and Fos-induction studies show that most sensory systems can activate neurons in the PBN (Campos et al., 2018, Carter et al., 2013, Kang et al., 2020), although the neuronal circuits involved are not established. Thus, the PBN is a hub activated by a wide variety of sensory signals, which then reports the state of the body to the other brain regions to elicit appropriate responses. All of these afferent signals to the PBN are excitatory (glutamatergic), but there are also inhibitory, GABAergic inputs including those from arcuate nucleus, bed nucleus of stria terminalis (BNST) and central nucleus of the amygdala (CEA). The PBN projects axons to the periaqueductal grey (PAG), extended amygdala (including the BNST, CEA, substantia innominata, SI), the cortex (primarily the insula), thalamus, parasubthalamic nucleus (PSTN), hypothalamus, and medulla.

Pioneering neuroanatomical studies encouraged functional studies (lesions, pharmacological and viral/genetic interventions), which have substantiated the predictions that the PBN is important for responding to internal and external stimuli and maintaining homeostasis. Examples include taste, thermal sensation, visceral malaise, pain, itch, hypercapnia, breathing, cardiovascular control, arousal, hunger/satiety, thirst, sodium appetite, and alarm (review, Palmiter, 2018). The response of the PBN to these sensory modalities raises multiple questions: How many different neuron populations are there? Are specific neurons or subsets of neurons involved in transmitting each signal? Is there integration of sensory signals (crosstalk between neurons) within the PBN? Do individual neurons project axons to one target region or send collaterals to many brain regions? A topographical map of the different neuronal populations in the PBN will help address these questions.

The PBN is bisected by a fiber tract, the superior cerebellar peduncle (scp) resulting in subregions that are lateral or medial to the scp in primates. In rodents, the scp is rotated relative to that in primates such that so-called lateral regions are more dorsal to the scp and medial regions are more ventral. The Kölliker-Fuse (KF) subregion is considered part of the PBN, while the cuneiform nucleus (CUN), mesencephalic trigeminal nucleus (MEV), and locus coeruleus (LC) are adjacent to it (Fulwiler and Saper, 1984, Paxinos and Franklin, 2019). *In situ* hybridization and immunohistochemistry studies have revealed that the PBN is primarily glutamatergic and expresses an abundance of different neuropeptides and neuropeptide receptors. These observations led to the creation of Cre-driver lines of transgenic mice that have been used to activate virally delivered, Cre-dependent genes to manipulate neuron activity and to visualize their axonal projections.

To define PBN cell types an gain insight unique expression of signaling molecules and receptors, we adopted the single-cell, RNA sequencing (scRNA-Seq) approach (Hashikawa et al., 2020, Macosko et al., 2015), which revealed 12 transcript-defined neuron types within the PB N proper ─ 90% of which are glutamatergic. We then used *in situ* hybridization to anatomically locate the neurons within the PBN and used Cre-driver lines of mice with viral expression of fluorescent proteins to establish the axonal projection patterns from many unique cell types.

## Results

### Single-cell RNA sequencing analysis of cell types in the PBN

To classify cell types in the mammalian PBN according to their transcriptional profiles, we harvested brain tissue from 10 adult male and female C57BL/6J mice, excised the PBN along its rostral-caudal extension (**Figure 1A**), and prepared cellular suspensions for high-throughput scRNA-Seq using a commercial droplet protocol (10X Genomics). After preprocessing the data to remove low-quality cells from the analysis (Rossi et al., 2021, Stuart et al., 2019), we retained a total of 39,649 single cell transcriptomes that were sequenced to a median depth of 47,177 reads per cell. A total of 17,038 genes were considered in the analysis, with a median of 1,740 expressed genes detected per cell **(Figure 1 – figure supplement 1)**.

**Figure 1.**
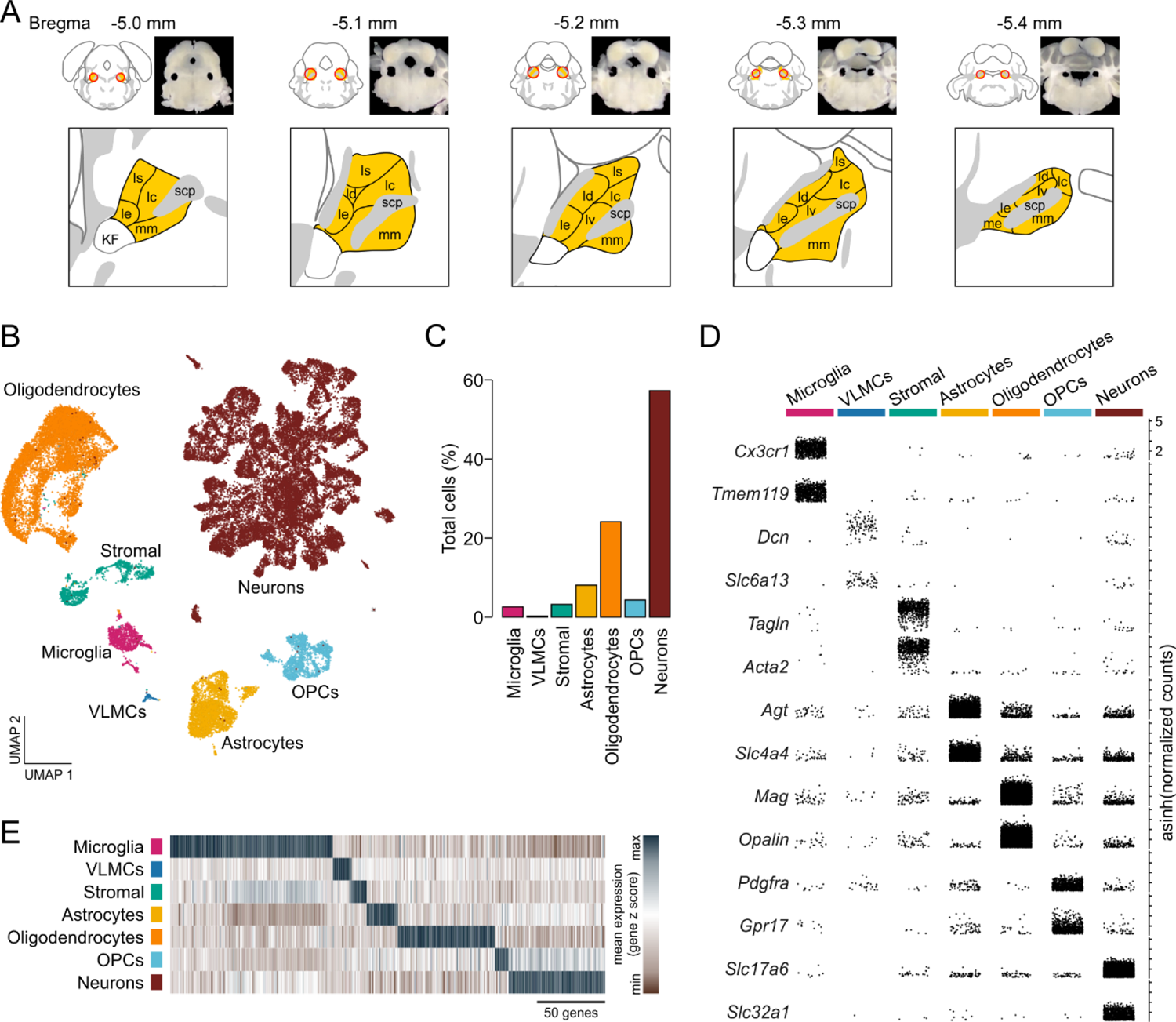
Single-cell RNA sequencing identifies resident cellular classes within the PBN. **(A)** Brain sections showing location of PBN and approximate boundaries of punches used for scRNA-Seq. PBN subregions from Allen Mouse Brain Atlas are shown in yellow; abbreviations are the same as in Figure 4. **(B)** Cells were clustered according to their transcriptional profiles and plotted in UMAP space. **(C)** Percentage of total cells comprised by each cluster. **(D)** Expression of canonical features across PBN clusters. Each point represents a single transcript plotted according to its asinh-normalized expression level. **(E)** Classes of PBN cell types are distinguished by unique transcriptional profiles comprised of multiple genes. **Supplementary File 1.** Table denoting the average normalized expression, fraction of cells expressing, and likelihood ratio p-value for every gene in each cluster.

To identify resident cell types of PBN tissue, cells were clustered on principal components and visualized in Uniform Manifold Approximation and Projection (UMAP) space (McInnes et al., 2018) (**Figure 1B**). We then applied a likelihood ratio test to identify features that were differentially expressed between clusters (Hafemeister and Satija, 2019, Macosko et al., 2015, McDavid et al., 2013), and classified cells according to the specificity of canonical marker genes within each cluster **(Methods, Figure 1B-E, and Supplementary File 1)**. Low resolution clustering identified 7 transcriptionally distinct populations of neurons, glia, and stromal cells within the PBN (**Figure 1B-E**). Of these, neurons represented the largest proportion of cells at 57.2% (**Figure 1C**). Oligodendrocytes, marked by *Mag* and *Opalin*, were detected at 24.1% of total cells and oligodendrocyte precursor cells (OPCs, 4.3% of total cells), were distinguished by their expression of *Pdgfra* and *Gpr17* (**Figure 1C-D**). The large number of oligodendrocytes and precursors is not surprising because the PBN is bisected by a large fiber tract, the superior cerebellar peduncle (scp). We detected a population of astrocytes representing 8.1% of cells that were labeled by robust expression of *Agt* and *Slc4a4*, as well as a smaller population of microglia (2.6%) that were specifically labeled by *Cx3cr1* and *Tmem119* (**Figure 1C-D**). Additionally, we identified two distinct populations of cells marked by stromal markers; one of these populations was characterized by its selective expression of *Tagln* and *Acta2* (3.2%), while another rare population of vascular leptomeningeal cells (VLMCs) was marked by *Dcn* and *Slc6a13* (0.26% of total cells). Although known canonical markers were used for the biased identification of broad classes of resident cells within PBN tissue (**Figure 1B and D**), each of these cellular classes was marked by a robust profile of unique transcriptional features (**Figure 1E and Supplementary File 1)**.

Next, we sought to identify distinct subclasses of neurons within the PBN. Subclasses of cells within a terminally differentiated cell-type share a similar transcriptional landscape, and as a result, statistically discriminating subclasses through clustering analysis requires the presence of a small set of high-confidence, high-variance features. To enable high-resolution subclustering of PBN neurons, we first applied a more stringent quality threshold to the neuronal population isolated in the initial analysis (**Figure 2 – figure supplement 1A-E)** resulting in a smaller set comprised of 7,635 neurons (**Figure 2A-B**). These cells were sequenced to a much higher median of 99,583 reads per cell with each cell containing a median of 3,189 genes and 7,823 transcripts **(Figure 2 – figure supplement 1F-O)**. We used a clustering approach like that applied for all cells and discriminated 21 unique subclusters of neurons (**N1-N21**) according to their expression of differential feature sets **(Methods, Figure 2,** and **Figure 2 – figure supplement 1P-Q)**. We first classified neuronal subclusters as GABAergic or glutamatergic; we designated **N1-N19** as glutamatergic with **N1, N2** being enriched in vesicular transporter *Slc17a7* (Vglut1), while **N3-N19** are enriched in vesicular transporter *Slc17a6* (Vglut2); the latter account for 90% of all neurons sequenced (**Figure 2C-E**). The remaining two subclusters **N20, 21** are GABAergic based on expression of the biosynthetic enzymes, *Gad1, Gad2*, and *Slc32a1* (Vgat)(**Figure 2C-E**).

**Figure 2.**
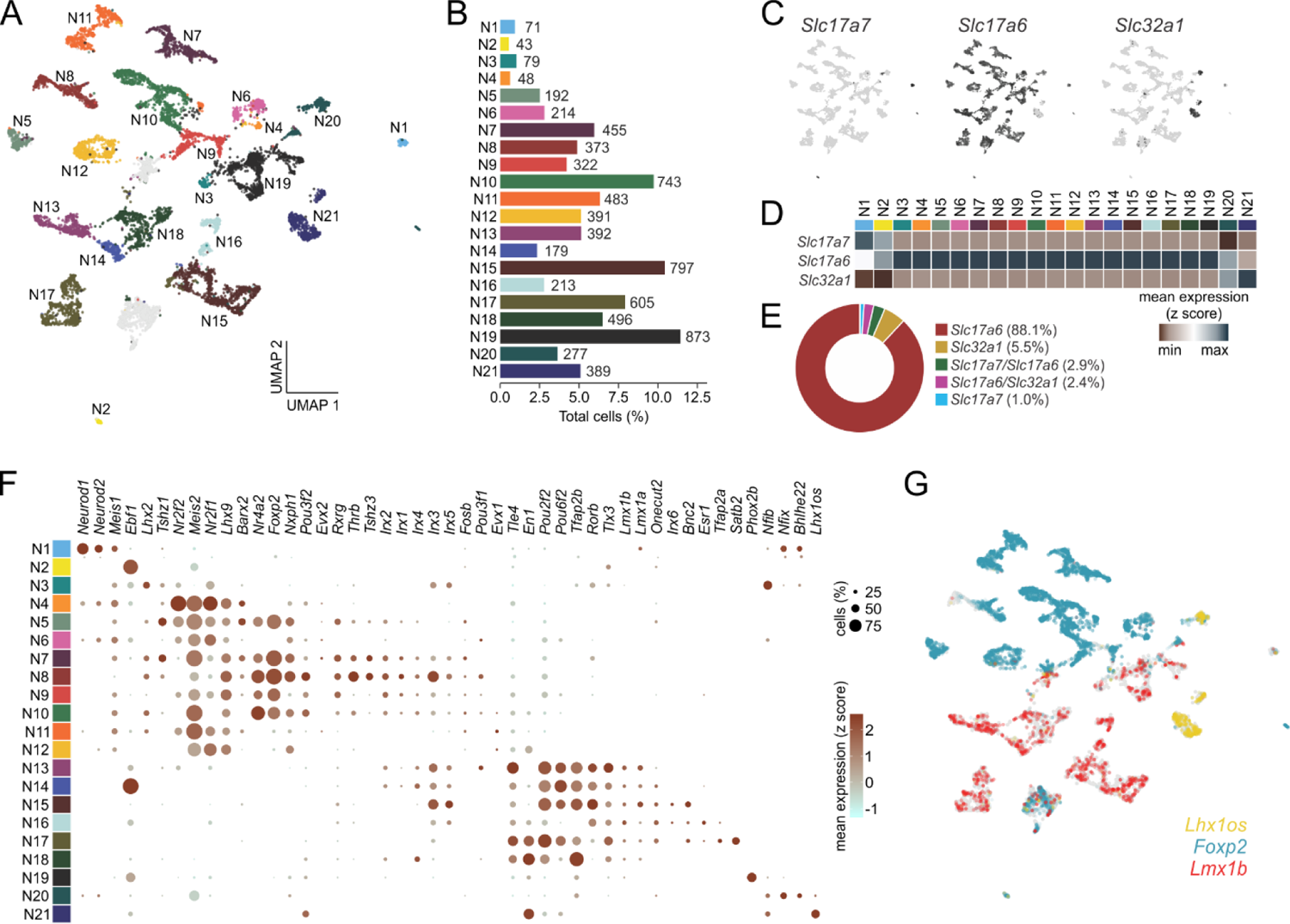
Single-cell RNA sequencing identifies discrete classes of PBN neurons. **(A)** Neurons were clustered according to their transcriptional profiles and plotted in UMAP space. **(B)** Percentage of total neurons comprised by each neuronal subcluster. **(C)** Expression values of fast neurotransmitters in UMAP space. **(D)** Average expression of fast neurotransmitters across neuronal subclusters. **(E)** Percentage of neurons individually expressing or co-expressing fast neurotransmitters. **(F)** Transcription factor expression across neuronal subclusters plotted according to their average normalized expression and fraction of cells expressing each gene. **(G)** Expression of the transcription factors *Foxp2*, *Lmx1b*, and *Lhx1os* across neurons in UMAP space. **(H)** Percentage of neurons individually expressing or co-expressing *Foxp2*, *Lmx1b*, and *Lhx1os*.

We sought to further define these 21 neuronal subclusters and identified three major clades that could be distinguished by their expression of transcription factors (**Figure 2F-G**). We found one major clade represented by *Lhx2, Lhx9, Meis2* and *Nrf1*, which are abundantly expressed in **N3-N12.** This clade probably descends from neurons that express *Atoh1* during development (Karthik et al., 2022). This group also includes nuclear receptors, *Nr2f1, Nr2f2, Nr4a*; zinc-finger protein, *Tshz1*; homeobox-containing transcription factors, *Barx2* and *Evx2*; and forkhead-box factor, *Foxp2*. *Nr4a2* and *Foxp2* represent a subgroup of this clade being expressed prominently in 5 of the 10 subclusters. The expression pattern of *Foxp2* in the PBN and its axonal projections via the ventral pathway to the hypothalamus have been described in detail (Huang et al., 2021a).

Another major clade includes **N15-N19** (**Figure 2F-G**) is represented by expression of a group of homeobox-containing transcription factors, *Pou2f2, Pou6f2, Lmx1a, Lmx1b, En1, Tlx3, Onecut2* along with a member of the retinoic acid family (*Rorb*), the AP2 factor (*Tfap2b*) and co-repressor (*Tle4*). *Satb2* (**N17**) and *Phox2b* (**N19**) represent individual subclusters within this clade.

The neuron subclusters that express *Slc17a7* (**N1 and N2**) and *Slc32a1* (**N20, 21**) are enriched in expression of basic helix-loop-helix proteins (*Neurod1, Neurod2, Bhlhe22*), *Nfib*, *Nfix* and *Lhx1os,* a long non-coding RNA transcribed from the *Lhx1* opposite strand (**Figure 2F-G**).

Within these three major clades, we also found that the 21 neuronal subclusters could be further delineated based on preferential expression of specific transcriptional features (**Figure 3A**). Although our analysis identified multiple known subclasses of PBN neurons, we also identified novel neuronal subclusters and their corresponding transcriptional markers. Thus, this analysis provides a representation of PBN neuronal diversity at a higher-resolution than has been previously appreciated (**Figure 3A-B**). A summary file listing the average expression of each gene per cluster, fraction of cells expressing each gene within a cluster, and the differential expression p-value per cluster is provided for all genes in the dataset **(Supplementary File 2)**.

**Figure 3.**
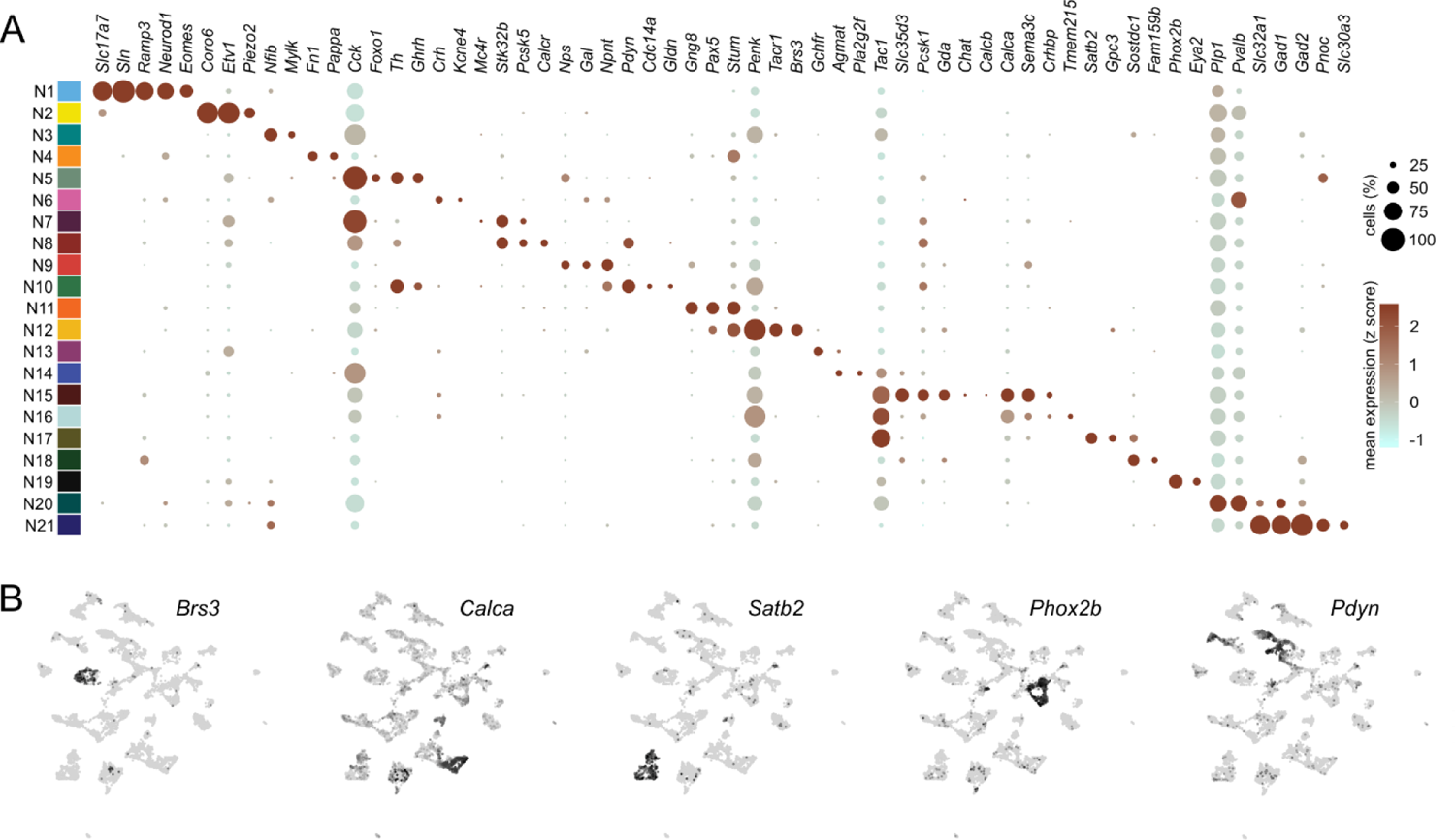
Distinguishing features of each neuronal subcluster. **(A)** Expression of select differentially expressed features across neuronal subclusters plotted according to their average normalized expression and fraction of cells expressing each gene. **(B)** Expression of select genes plotted in UMAP space. Figure 3 **– table supplement 1.** Neuropeptides and GPCRs with restricted expression. **Supplementary File 2.** Table denoting the average normalized expression, fraction of cells expressing, and likelihood ratio p-value for every gene in each neuronal subcluster. **Supplementary File 3.** Expression of neuropeptides across neuronal subclusters plotted according to their average normalized expression and fraction of cells expressing each gene. **Supplementary File 4.** Expression of GPCRs across neuronal subclusters plotted according to their average normalized expression and fraction of cells expressing each gene.

### Expression of neuropeptides and G-protein coupled receptors in the PBN

Neuropeptides are valuable markers for selected subpopulations of neurons in many brain regions. Some of these neuropeptide mRNAs are uniquely and robustly expressed in one or two subclusters, e.g., *Calca, Ghrh, Nps, Npy, Pdyn*, *Pnoc*, while others are expressed in multiple subclusters, e.g., *Adcyap1, Cck, Crh, Gal, Gpr, Nmb, Nts, Penk, Tac1, Vgf* (**Figure 3 – table supplement 1, Supplementary File 3**). Like the neuropeptide group, the genes encoding G-protein-couped receptors (GPCRs) are genetically useful because they are likely to be expressed in neurons that receive aminergic or neuropeptide input; thus, making them desirable for circuit mapping **(Supplementary File 4)**. Indeed, Cre-driver lines of mice have been generated for many GPCR genes. GPCRs are also of interest because they are viable targets for drugs that could potentially modify the function of select neurons and circuitry in the PBN. The expression of GPCR genes is generally much lower than that of neuropeptides, making them difficult to detect by immunohistochemistry or *in situ* hybridization. Consequently, scRNA-Seq data provide a much-needed resource for identifying the GPCRs expressed by distinct PBN neurons. GPCRs with relatively restricted expression are listed in **Figure 3 – table supplement 1.**

### Locating neuronal populations by *in situ* hybridization

To determine if different neuron subclusters are located in distinct PBN subdivisions, we performed fluorescent *in situ* hybridization on coronal sections of the PBN spanning the rostral-caudal axis from Bregma −5.0 to −5.4 mm. Representative probes for each neuronal subcluster (**N1 to N21**) were chosen based on the distinguishing genes in each subcluster (**Figure 4A**). A composite image of each PBN section was generated and PBN subdivisions were drawn using a combination of probe expression and the Allen Mouse Brain Atlas (AMBA) as a guide (**Figure 4B**). We localized several neuron subclusters within major AMBA subdivisions of the PBN (**Figure 4C** and **Figure 4 – figure supplement 1)** and qualitatively scored the expression of each probe within a specific PBN subregion (**Figure 5A**) or neighboring regions **(Figure 5 – figure supplement 1)** based on the number of transcripts (signal intensity) and number of cells (**Figure 5B**). The discussion below is derived from **Supplementary File 5**, which includes results for all HiPlex probes at 5 Bregma levels throughout the PBN. RNAscope experiments were also used to resolve co-expression issues as indicated below.

**Figure 4.**
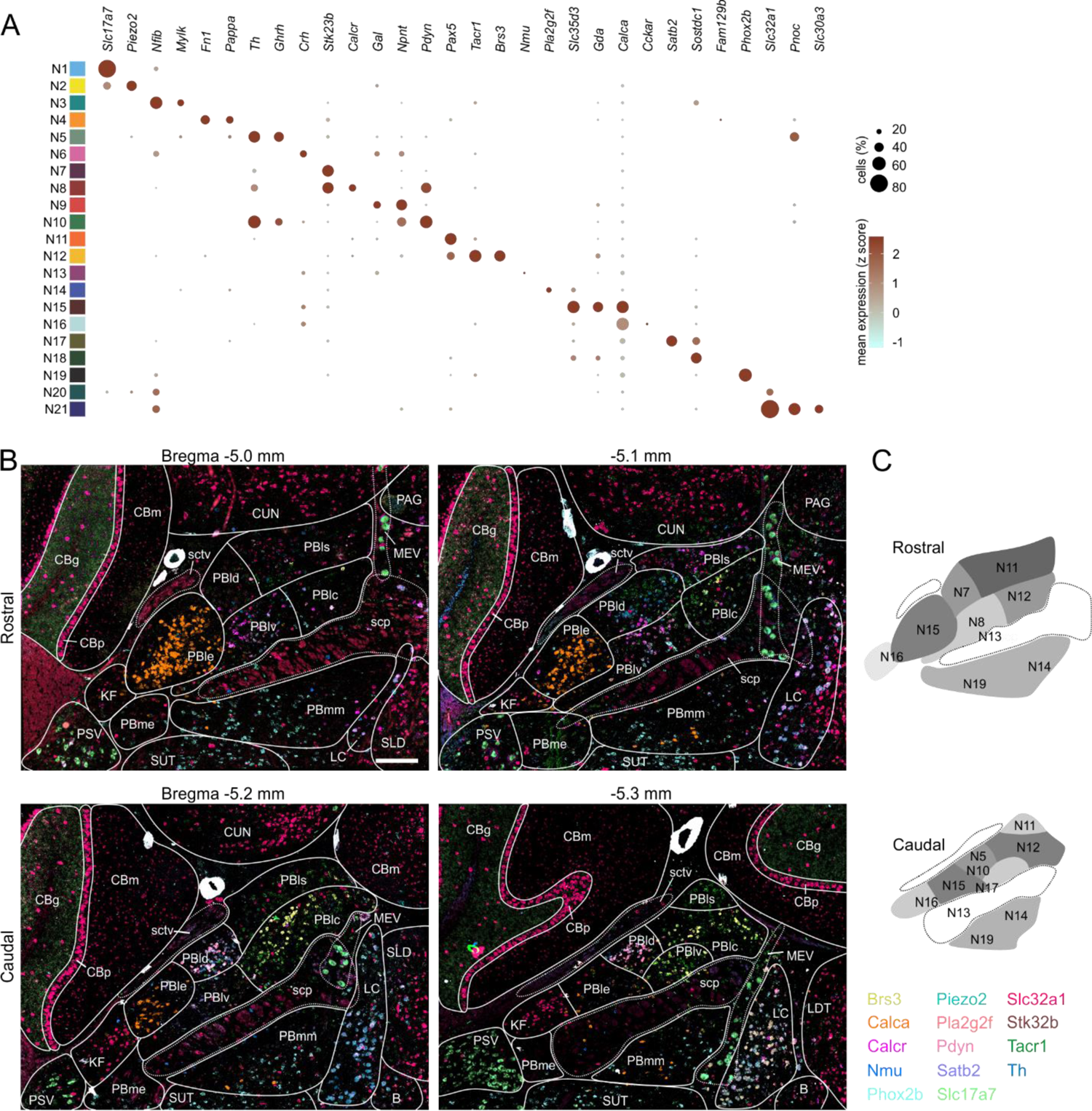
Localization of mRNAs in subregions of the PBN based on HiPlex results. **(A)** Expression of genes selected as HiPlex probes within the scRNA-seq dataset. **(B)** Example of how the ROIs denoting PBN subregions were drawn for analysis of 15 probes from the first HiPlex experiment. Probes and their colors are indicated. **(C)** Diagram of the approximate location of 12 of the identified clusters in a rostral and caudal PBN. Scale bar, 200 μm. Abbreviations from AMBA: PBle, external lateral PBN; PBlv, ventral lateral PBN; PBld, dorsal lateral PBN; PBlc, central lateral PBN; PBls, superior lateral PBN; PBmm, medial PBN; PBme, medial external PBN; scp, superior cerebellar peduncle; KF, Koelliker-Fuse; sctv, ventral spinocerebellar tract; CUN, cuneiform nucleus; LDT, laterodorsal tegmental nucleus; MEV, Midbrain trigeminal nucleus; SUT, supratrigeminal nucleus; SLD, sublaterodorsal nucleus; B, Barrington’s nucleus; PSV, principle sensory trigeminal nucleus; CBm, cerebellar molecular layer; CBg, cerebellar granular layer; CBp, cerebellar Purkinje layer; PAG, periaqueductal gray; LC, locus coeruleus. Figure 4 **– figure supplement 1.** Example of HiPlex staining for *Brs3*, *Calca*, and *Phox2b* for 5 Bregma levels. Solid lines surround clusters of positive neurons or individual neurons, colored dashed lines indicate expression outside the PBN, such as in KF and LC. **Supplementary File 5**. HiPlex data for all probes at 5 Bregma levels of the PBN.

**Figure 5.**
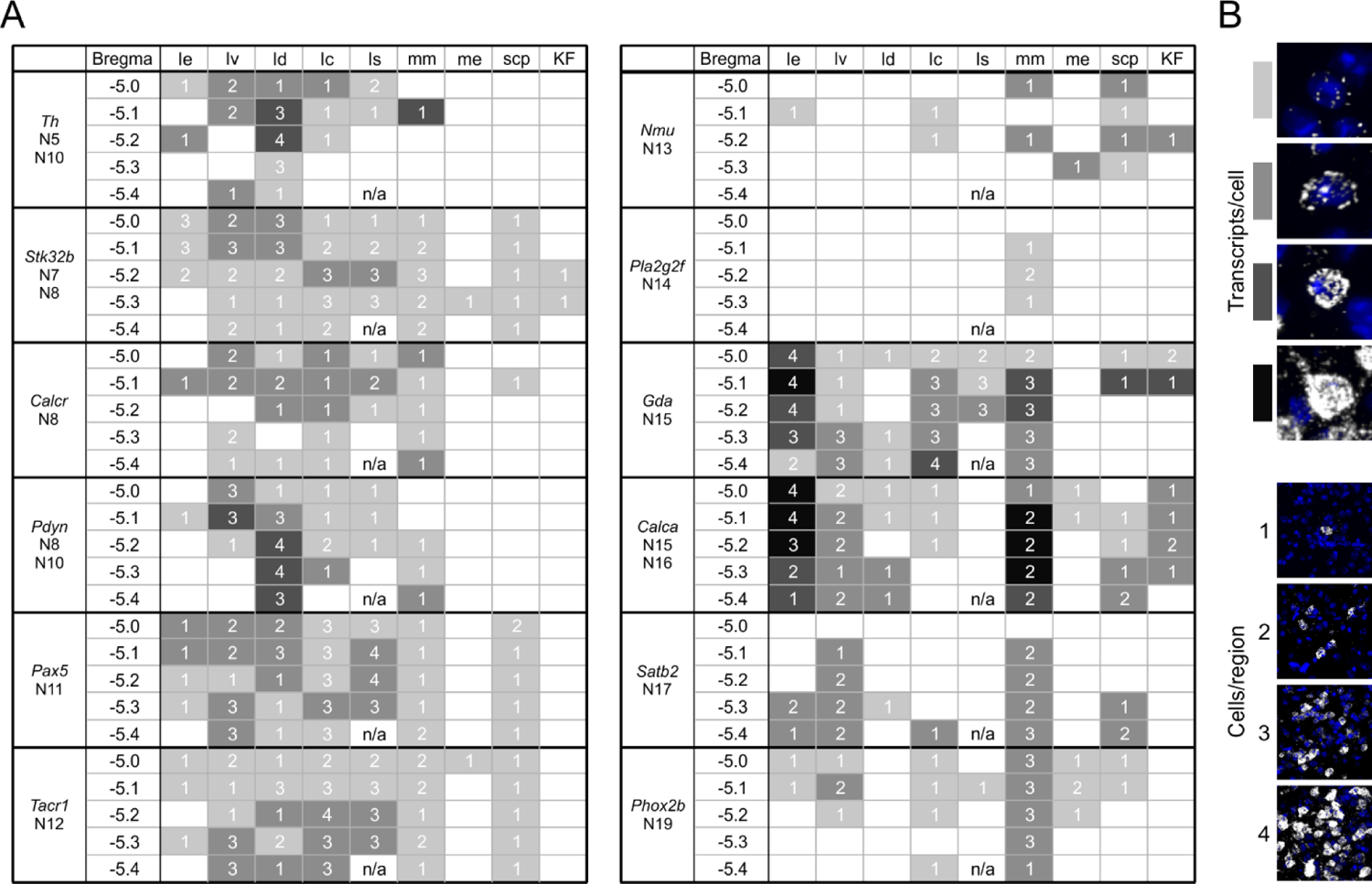
A guide to HiPlex results showing relative strength and abundance of mRNA expression in PBN subregions. **(A)** Qualitative expression for 12 genes that can be used to identify different subregions in the PBN. Strength of expression and percentage of cells in subregions of the cells were analyzed for 5 bregma levels. **(B)** Key for the colors and numbers in the table. Shade of gray gets darker as the number of transcripts per cell increases and the number represents an estimate of the number of positive cells per subregion. The abbreviations are defined in Figure 4. **Figure 5 – figure supplement 2.** *Pdyn* and *Calca* neurons form tight clusters in PBld and PBle, respectively. **Figure 5 – figure supplement 3.** *Calca/Gda or Calca/Tac1/Satb2* co-expression analyzed by RNAScope **Figure 5 – table supplement 4.** Table showing enrichment of neuropeptide mRNAs in *Calca* neurons based on RiboTag experiment **Supplementary File 6.** Source data for Ribotag experiment showing all genes.

#### N1 and N2

*Slc17a7* (Vglut 1) and the transcription factors, *Neurod1*, *Neurod2* and *Meis1* are expressed robustly in the Cb granular layer; some of these non-PBN cells were included in the tissue punch and probably represent **N1**; however, there are also few *Slc17a7*-positive neurons scattered throughout the PBN, especially in PBld. *Piezo2, a* marker for **N2**, is robustly co-expressed with *Slc17a7* in mesencephalic trigeminal neurons (MEV). Note that both of these clusters represent <2% of PBN neurons (**Figure 2B**).

#### N3, N4

*Mylk* and *Fn1* were chosen to represent **N3** and **N4**, respectively; however, *Fn1,* and to a lesser extent *Mylk*, had weak signals throughout the sections in what appear to be blood vessels. In a repeat experiment, they were replaced with *Nfib* (**N3**) and *Pappa* (**N4**). *Nfib* probe had a weak signal in several regions outside the PBN (PSV, CB and LDT). *Pappa* was also expressed outside the PBN (MEV, B, CB) with weak expression in caudal PBld sections that overlap with *Pdyn, Crh,* and *Pnoc* (**N10)**. These clusters also represent <2% of PBN neurons (**Figure 2B**).

#### N5

*Th, Ghrh* and *Pnoc* were chosen to represent **N5** although they are not uniquely expressed in **N5**. There is a group of *Th*-positive cells in rostral PBld that do not express *Pdyn* that could be **N5**; however, they do not express *Ghrh* or *Pnoc* as predicted by scRNA-Seq. TH protein expression in lateral PBN was documented (Milner et al., 1986) and *Th* mRNA is expressed in lateral PBN based on AMBA; it is also expressed in LC, as expected. *Pnoc* expression is scattered throughout PBN and neighboring nuclei and its expression overlaps with *Slc32a1* in some GABAergic cells (**N21**).

#### N6

*Crh, Npnt* and *Gal* were chosen to locate **N6**. *Crh and Npnt* are co-expressed in many cells in the sublaterodorsal nucleus (SLD) that is adjacent to the PBN in rostral sections, but *Gal* is not co-expressed with *Crh* in SLD. *Crh* is also expressed in PBle, where it is co-expressed with *Calca (***N15,16**), as well as in PBlv, PBld, B and LC regions.

#### N7

*Stk32b* was chosen to localize **N7**. It is expressed widely throughout the PBN and neighboring brain regions. Its expression overlaps with other markers, e.g., some *Calca* and *Pdyn* neurons; however, there is a *Stk32b* cluster in rostral PBld that does not overlap with other markers that could represent **N7**. *Cck* and *Nps* are predicted to be robustly expressed in **N5** and **N7** (**Figure 5A**) but they were not chosen for HiPlex; *Cck* is widely expressed in other clusters as well.

#### N8, N9, N10

scRNA-Seq clustering analysis suggested that *Pdyn* would be co-expressed with *Stk32b* and *Calcr* in **N8**, which was confirmed by HiPlex analysis. This cluster is closer to the scp in the rostral PBlv region. In addition, **N10** is represented by *Pdyn, Ghrh, Pnoc, Npnt* and *Th*, which was confirmed by HiPlex; these neurons are in more caudal PBld regions. *Npnt* without *Pdyn* was chosen to represent **N9**, but the few cells of this type also express *Crh* (**N6**); thus, the location of **N9** is uncertain. Although tyrosine hydroxylase (*Th*) mRNA is expressed in many *Pdyn* neurons, it is unlikely that they are catecholaminergic because the protein levels are very low compared to LC (Karthik et al., 2022) and other genes required for catecholamine synthesis, vesicular transport and re-uptake were not detected. Transduction of *Pdyn^Cre^* mice with AAV carrying Cre-dependent effector genes has been used to assess their functions, which include temperature regulation, nocifensive behaviors and feeding (Chiang et al., 2020, Geerling et al., 2016, Kim et al., 2020, Norris et al., 2021, Yang et al., 2021). It will be important to learn whether the two clusters of *Pdyn* neurons have distinct functions.

#### N11, N12

The transcription factor *Pax5* was chosen as a marker for **N11** and **N12**. There is a distinct group of *Pax5*-positive cells in the rostral PBlv close to scp that does not overlap with *Pdyn* that could represent **N11***. Brs3* and *Tacr1* are co-expressed and represent a subset of *Pax5* neurons in rostral PBlc and PBls as **N12**. Expression of *Tacr1* and *Pax5* and expression is more widespread than *Brs3,* extending into PBls and PBmm in caudal sections. The *Pax5* signal is more diffuse than other probes as if it is not restricted to the cell body. Studies using *Tacr1^Cre^* mice revealed that they play important roles in nocifensive behaviors (Barik et al., 2021, Deng et al., 2020, Huang et al., 2019).

#### N13, N14

*Nmu* was chosen for **N13**. A small number of *Nmu*-expressing neurons was observed in the mid-PBN sections that are scattered within the scp or adjacent PBmm, in agreement with location of cell bodies after injecting *Nmu^Cre^*mice with a Cre-dependent fluorescent reporter (Jarvie and Knight, personal communication). *Pla2g2f* was chosen to locate **N14**; there is a small cluster of weakly expressing cells in the PBmm of mid-PBN sections. Note that the transcription factor *Ebf1* is robustly expressed in **N14** and that this cluster is represented by only ∼2.5% of neurons (**Figure 2B, F**).

#### N15, N16

*Calca* is a defining gene of PBle as revealed by AMBA, our Hiplex analysis, immunohistochemistry (Shimada et al., 1985), viral expression of reporter genes activated by injection into the PBN of *Calca^Cre^* mice (Bowen et al., 2020, Chen et al., 2018, Huang et al., 2021b, Kaur et al., 2017). *Calca^Cre^* mice have been used extensively to examine the role of these neurons in nocifensive behaviors, appetite, and arousal (Palmiter, 2018). It was unexpected that the scRNA-Seq analysis would reveal *Calca* expression in 2 clusters. **N15** neurons are 3-fold more abundant than **N16** neurons (**Figure 2B**). *Calca* is expressed robustly in PBle, but a few *Calca*-expressing cells are scattered throughout the lateral and medial PBN, which is especially prominent in rostral sections. *Calca* expression extends into the KF where it is intermingled with *Slc32a1*-expressing neurons. It is also expressed in the LC, especially in caudal sections. To distinguish between the two clusters, we used RNAScope with probes for *Calca* and *Gda* (**N15** enriched). *Calca* cells with strong expression in PBle almost entirely overlap with *Gda* along with both strongly and weakly expressing Calca cells in other sub-regions, indicating **N15.** There are *Calca* cells without *Gda* scattered sparsely throughout the PBN and there is a small group of *Calca* cells that do not express *Gda* in the lateral ventral PBle that extend partially into the KF that likely represent **N16**. (**Figure 5 – figure supplement 2**).

Most of the *Calca*-expressing neurons within PBle form a tight cluster without other neurons interspersed. RNAscope *in situ* hybridization with probes for *Calca* and *Slc17a6* reveal co-expression with no *Slc17a6*-only signal within the core of PBle, but there is some intermixing of neurons bordering the PBle (**Figure 5 – figure supplement 3**). This arrangement was also more apparent when *Slc17a6* expression was inactivated using a virus expressing Cas9 and two guide RNAs targeted to *Slc17a6*; in that case the *Slc17a6* signal was uniformly weak in the PBle. Importantly, no *Slc17a6*-expressing cells were interspersed among *Calca*-expressing neurons (data not shown). There are no GABAergic neurons in the core of PBle either (compare *Calca* and *Slc32a1* in **Supplementary File 5**).

Both the *Calca* and *Calcb* genes encode calcitonin gene-related peptide (CGRP), but *Calcb* is only weakly expressed in cluster N**15** based on scRNA-Seq, although the AMBA shows robust expression of *Calcb* perhaps because their probe hybridized to both genes. The weak expression of *Calcb* was confirmed by immunohistochemistry experiments since *Calca*-null mice express negligible CGRP in the PBN (Chen et al., 2018, Zajdel et al., 2021).

The co-expression of several neuropeptides along with CGRP was verified by immunoprecipitation of polysomes from *Calca^Cre^* mice expressing *Rpl22^HA^* (RiboTag) followed by microarray analysis of mRNAs (Sanz et al., 2009), which revealed enrichment of *Crh, Nts, Tac1, Adcyap1* and *Vgf* (**Figure 5 – table supplement 5**); these results were confirmed by scRNA-Seq, which also identified several more neuropeptides expressed in these neurons **(Supplementary File 3)**. Expression of two GPCR mRNAs (*Avpr1a* and *Galr1*) is restricted to **N15** and *Cckar* is restricted to **N16 (Figure 3 – table supplement 1)**. Several other GPCRs, including *Oprm1* which plays an important role in opioid-induced respiratory depression (Liu et al., 2021), are expressed along with *Calca*, but also in other clusters (**Supplementary File 4)**. Unexpectedly, *Chat* (encodes acetylcholine biosynthetic enzyme) is expressed in **N15** (as well as **N6**), a result consistent with location fluorescent protein expression after viral transduction of *Chat^Cre^* mice with AAV-DIO-YFP (see below) and immunohistochemistry (Garfield et al., 2015). *Slc18a3,* the vesicular transporter for acetylcholine, is selectively expressed in **N15**, suggesting that this *Calca* neuron population is both glutamatergic and cholinergic.

#### N17

*Satb2*, which encodes a transcription factor, is a defining gene for this cluster. These neurons are scattered in the PBlv, and PBmm and scp in the caudal PBN based on Hi-Plex results and reporter gene expression from *Satb2^Cre^* mice. RNA-Seq experiments predicted that *Satb2* neurons co-express *Tac1,* which was confirmed by an RNAScope experiment in which *Tac1, Satb2* and *Calca* probes were combined (**Figure 4 – figure supplement 4**). This experiment revealed co-expression of *Tac1* and *Calca* as well as *Tac1* and *Satb2*, but there were also abundant *Tac1* cells without expression of either *Satb2* or *Calca*, especially in PBlv and PBlc. *Satb2^Cre^* mice have been used to show that these PBN neurons relay taste signals to the thalamus (Fu et al., 2019, Jarvie et al., 2021).

#### N18

*Shisal2b* and *Sostdc1* were chosen to represent **N18**. Weakly expressing *Sostdc1* cells were scattered throughout caudal PBN sections. No *Shisal2b* signal was detected in any section examined.

#### N19

The transcription factor *Phox2b* was chosen as a distinct marker for **N19**; there is sparse expression *of Phox2b* in PBmm within and around the Scp. There is more abundant expression of *Phox2b* on the medial side of the PBN and extending into the neighboring LC and KF in agreement with (Karthik et al., 2022).

#### N20, N21

The vesicular transporter for GABA (Vgat) encoded by the *Slc32a1* gene is expressed along with the GABA biosynthetic enzymes, *Gad1* and *Gad2*, in neurons scattered throughout the PBN, with some small clusters of *Slc32a1*-positive neurons in PBls, PBmm and KF. GABA neurons within the PBN can inhibit local glutamatergic neurons (Sun et al., 2020). RNA-Seq analysis indicated that *Pnoc* is expressed in **N21** but not **N20**. Most *Slc32a1*-expressing cells in KF do not express *Pnoc*. Some GABAergic cells scattered throughout the rest of the PBN express *Pnoc* while others do not; thus, **N20** and **N21** appear to be intermingled throughout the PBN and KF. In summary, of the neuron subclusters identified by scRNA-Seq, some are in brain regions that are adjacent to the PBN including **N1, 2, 3, 4 and 6**; some clusters, e.g., the GABAergic clusters, are sparse and scattered throughout the PBN (**N9, 18, 20, 21**); leaving twelve clusters (**N5, 7-8, 10-17, 19**) that can be placed into one of the AMBA sub-regions (**Figure 4C**).

### Mapping PBN expression and axonal projections with Cre-driver lines of mice

Expression of fluorescent proteins from AAV injected into the PBN of Cre-driver lines of mice provides an independent means of locating the cell bodies; it also allows one to visualize the axonal projections. We injected AAV1-Ef1a-DIO-YFP and AAV1-Ef1a-DIO-synaptophysin:mCherry into the PBN or surrounding region of 21 Cre-driver lines to visualize cell bodies and processes within the PBN (YFP) and synapses (mCherry) throughout the entire brain. Cre-expressing cells in the PBN were categorized by their location within large PBN subdivisions and surrounding regions (**Figure 6A-E**). In addition to the 21 Cre-driver lines described here, PBN expression from additional Cre-driver lines has been reported (**Figure 6, table supplement 1**), not including those described in the Allen Institute Connectome project. *Slc17a6^Cre^* (Vglut2), which is expressed in most of the clusters, reveals the overall distribution of glutamatergic projections from the PBN (Huang et al., 2019). As expected, the location(s) of fluorescent cell bodies in the PBN of the Cre-driver lines of mice is consistent with *in situ* hybridization results, although the promoter in the virus and multiple viral particles per cell can provide more robust expression than the endogenous gene, e.g., many GPCRs where the *in situ* signal in the AMBA is undetectable.

**Figure 6.**
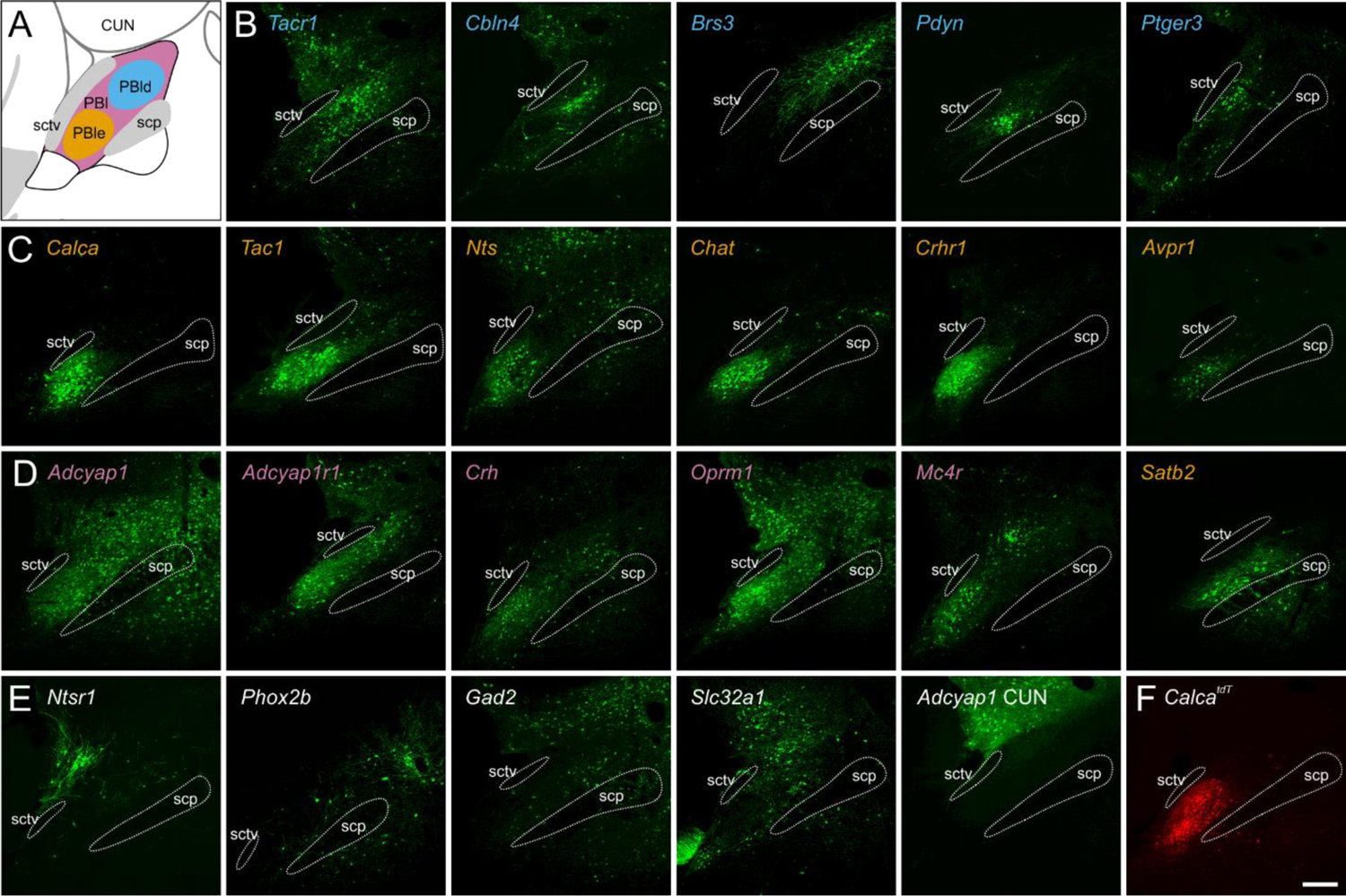
PBN expression in Cre-driver mouse lines. **(A)** Schematic of the PBN showing PBle in orange dotted line, dorsal PBN regions in blue dotted line, and expression in both in pink. **(B**) Five Cre-driver lines (blue lettering) with expression primarily in dorsal PBN**. (C)** Six Cre-driver lines (orange) primarily in PBle. *Satb2* is included here because its projection pattern resembles that of this group. **(D)** Five Cre-driver lines (pink) with expression in several PBN regions. **(E)** Five Cre-driver lines (grey) with expression patterns that do not fit with the other categories. **(F)** Image of *Calca-tdTomato* expression in the PBN for comparison. All images of viral expression are in mid-PBN sections; approximately Bregma −5.2 mm. Scale bar, 200 μm. Figure 6. Source data available at Zenodo DOI: 10.5281/zenodo.6707404 and includes complete TIFF stacks for each of these Cre-drivers and *Calca-tdTomato*.

We also mapped the expression of *Calca* in mice with tdTomato targeted to the *Calca* locus (Calca^tdT^, **Figure 6F).** Fluorescence from *Calca^tdT^* mice reveals that in addition to robust expression in PBle, a few *Calca*-expressing cells are scattered throughout the lateral and medial PBN, which is especially prominent in rostral sections in agreement with the *in situ* experiments above. This approach also reveals tdTomato fluorescence that extends into the KF. The LC, trigeminal and facial nuclei also express Calca, in agreement with AMBA. Note that a genetic cross of *Calca^Cre^* mice with Cre-dependent reporter line, e.g., Gt(*ROSA)26Sor^lsl-tdTomato^* (Ai14) results in widespread fluorescence throughout the brain due to developmental expression (Carter et al., 2013).

For each of the 21 Cre-driver lines that we analyzed and the *Calca^tdT^* line, images were taken from every 3^rd^ coronal, 35-μm section, stitched together and registered to generate a TIFF stack that can be manipulated to view expression of YFP and mCherry throughout the brain using ImageJ. All images are available to view and download from Zenodo (DOI: 10.5281/zenodo.6707404); an example video of *Calca* cell expression is included here (**Video 1**).

Cell populations in the PBN can conceptually be divided into two major groups based on overall projection patterns (**Figure 7A**). The relatively dense cluster of cells in the PBle generally follows the central tegmental tract (CTT) and the cells in the dorsal regions (PBld/ls/lc) tend to follow the ventral pathway (VP). Both populations also project along the descending pathway into the brainstem, but the projections to the brainstem from the cells in the dorsal regions tend to be weaker than those from the PBle (**Figure 7B**). Cells that reside in the PBmm do not appear to have a unique projection pathway, but rather offer variations in projection strength along the two major pathways.

**Figure 7.**
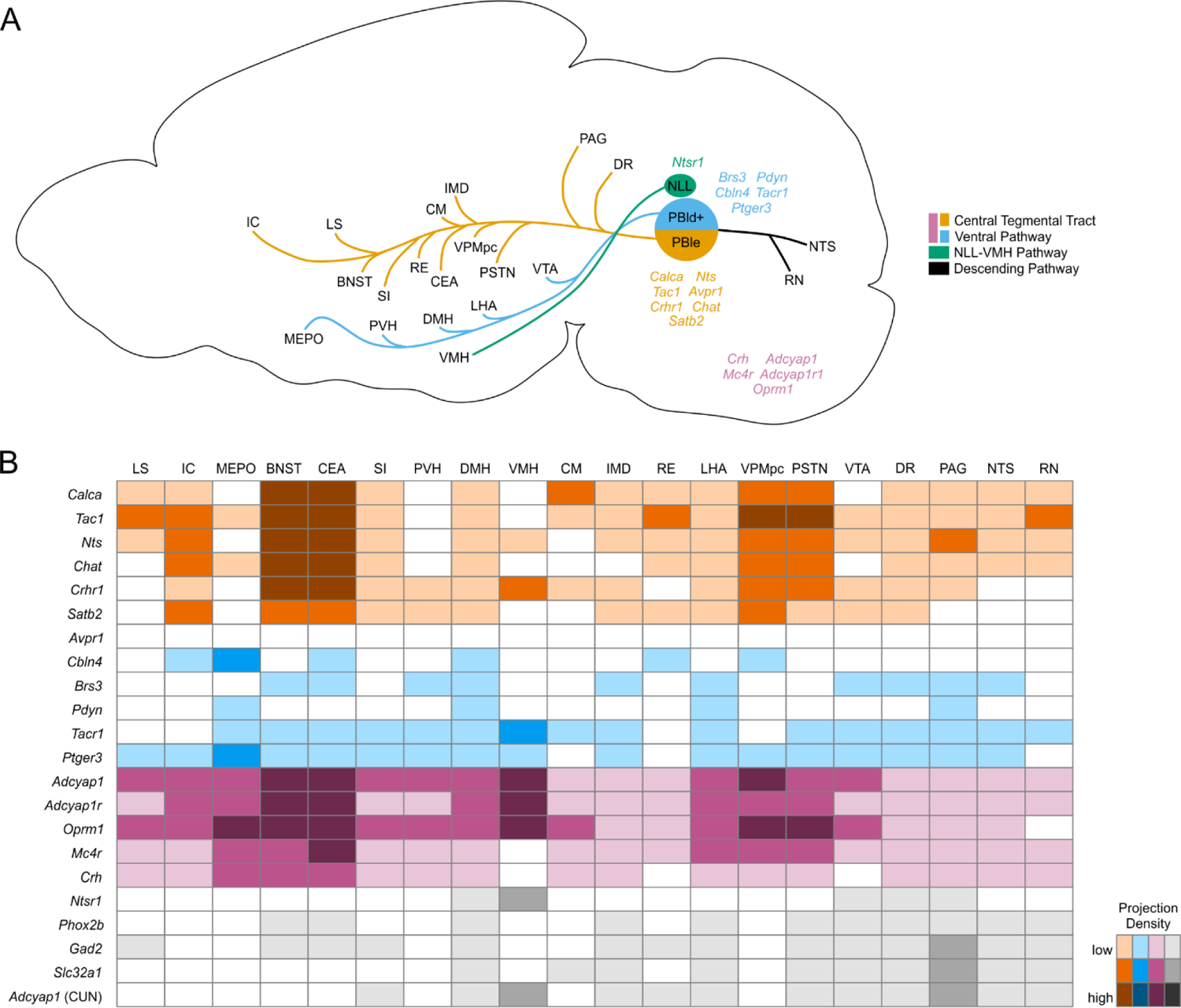
Descending pathways from the PBN and surrounding regions and the strength of their projections. **(A)** Diagram showing the two main ascending projection pathways from the PBN. Genes are listed in a matched color with their pathway; pink genes follow both pathways. Many genes from each group have projections into the descending pathway that are not shown. (**B)** Guide showing approximate density of synaptophysin in a subset of target regions along with their abbreviations. Colors represent the pathways; darker shades indicate denser innervation.

For Cre-driver lines that are expressed almost exclusively in the PBle (*Calca, Crhr1, Nts, Chat*), or strongly in the PBle along with other regions (*Crh, Adcyap1, Adcyap1r1, Tac1, Mc4r, Oprm1*) the axon terminals with the brightest signal are in the BNST and CEA. Although expression is generally spread throughout these subregions, the oval and ventral regions of the BNST and the capsular region of the CEA have the strongest fluorescence. It is likely that the PBle cells are responsible for most of the oval/ventral BNST and capsular CEA staining density, while other neurons in the dorsal PBN project to adjacent regions. Although there is some variation, the next brightest regions are the IC, DMH, SI, VPMpc, and PSTN. There are no lines that have expression in the PBle that do not have at least some amount of synaptophysin in the above listed regions. Other areas that are often innervated are the LS, CM, LHA, RE, IMD, PAG, DR, RN, and NTS. Many of the Cre-driver lines that have most of their expression in PBle still have cells scattered throughout other subregions in the PBN, which may explain the expression in areas that have been associated with the ventral tract, such as the DMH and LHA.

Cre-driver lines that mainly have expression in the dorsal regions (*Pdyn, Tacr1, Brs3, Cbln4, Ptger3*) tend to target ventral brain regions such as the MEPO and DMH. The cell groups in the dorsal areas do not always overlap with the PBle groups; the intensity of projection sites can vary in areas like the MEPO from very strong (*Cbln4*) to very weak (*Pdyn*). Cre-driver lines with smaller cell populations (*Pdyn, Brs3*) innervate fewer regions with weaker signal such as PVH, LHA, DMH, and PAG, than lines with more cells (*Tacr1, Ptger3*). Cre-driver lines that include some expression in PBle weakly innervate areas associated with the CTT such as BNST and CEA.

Some Cre-driver lines have strong cellular expression across most of the lateral PBN. For lines like this (*Adcyap1, Adcyap1r1, Oprm1, Crh*), there is often strong innervation in areas associated with the CTT such as the BNST/CEA and areas associated with the VP such as the MEPO. Overall, their projections are a combination of areas seen in the other groups.

The cells that reside in the PBmm do not appear to have a separate innervation profile. There was no line examined that had its expression limited to the PBmm exclusively. However, in one example (*Tac1*), the cells were transduced in PBmm on only one side, allowing for a comparison of projections between hemispheres. The result was a slightly brighter synaptophysin signal in the common projection targets of IC, BNST, and CEA, and a much brighter signal in the cortical amygdala on the side with the PBmm expression. In another example (*Phox2b*), most of the cells transduced were in the PBmm, with some cells in the PBlc and PBls. This line showed a unique projection to the SH, which likely originated from cells in the PBlc and PBls rather than PBmm because we did not see the same SH projection from the *Tac1* cells in the PBmm (*Phox2b* and *Tac1* whole brain expression available on Zenodo, DOI: 10.5281/zenodo.6707404).

A few Cre-driver lines reveal unique projection patterns. *Satb2* is a population that resides mainly in PBlv, scp, and PBmm. Although only partially expressed in PBle, the projection pathway still largely follows the CTT with connections to areas typically associated with PBle such as BNST and CEA, but with weaker projections to the oval and capsular subregions typical of PBle lines. The axon terminals from *Satb2^Cre^-* and *Calca*^Cre^-driver lines in the BNST and CEA reveal that they are distinct (**Figure 8A-B**). *Ntsr1* is another population that does not match the others. Its cell bodies are dorsal to the rostral PBN in the nucleus of the lateral lemniscus (NLL). The main projection target is the VMH (**Figure 8C**). All other lines that have projections to the VMH (*Crhr1, Nts, Adcyap1, Adcyap1r1, Tacr1, Oprm1*) also have cell bodies in the same region as *Ntsr1*, which suggests that projections to VMH likely come from viral transduction of cells in the NLL rather than the PBN.

**Figure 8.**
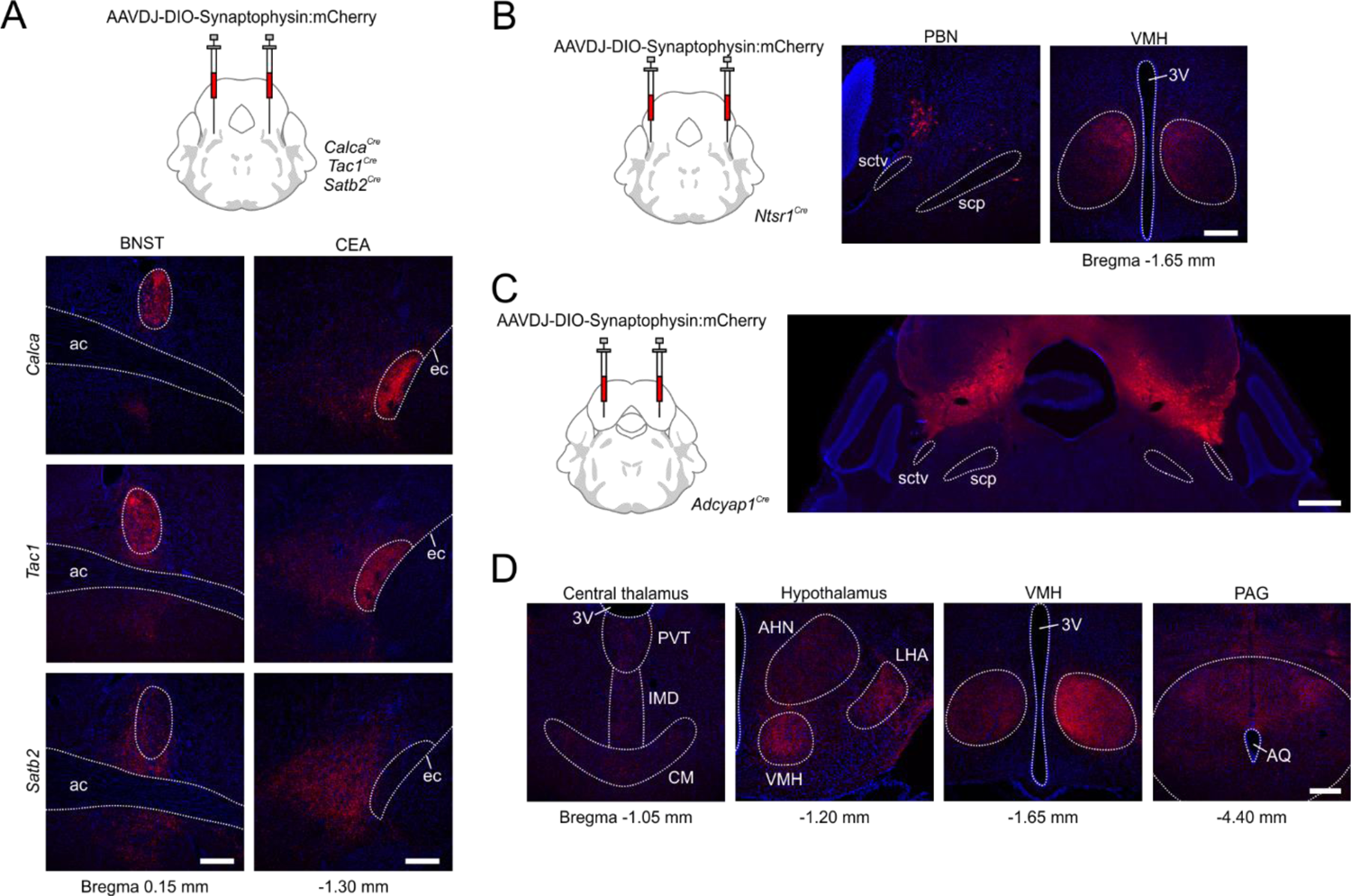
Projection patterns in target brain regions. (**A)** Comparison of synaptophysin:mCherry staining in BNST and CEA for *Calca, Tac1* and *Satb2* Cre-driver lines. Scale bars, 200 μm. (**B)** Example of cell body location in NLL region adjacent to PBN for *Ntsr1^Cre^* mice (left); they project almost exclusively to the VMH (right). Scale bar, 200 μm. **(C)** Cell bodies in the CUN, ICE, and PAG, but not PBN, in *Adycap1^Cre^* mice. Scale bar, 500 μm. **(D)** Examples of projections from *Adycap1*-expressing neurons in non-PBN regions that may mistakenly be attributed to the PBN. Scale bar, 200 μm.

Many of the genes of interest are widely expressed in regions surrounding the PBN, which made restricting the transduction of cells within the PBN nearly impossible. *Gad2* and *Slc32a1*, the two GABAergic genes, are only sparsely expressed within the PBN compared to adjacent areas; consequently, we were unable to determine whether they function as interneurons or as projection neurons. *Gad2* is expressed more widely and without *Slc32a1* in PBN glutamatergic cells, e.g., *Calca* neurons, so using that Cre-driver line to examine inhibitory projections outside of the PBN could be misleading. To explore the projection pathways that may arise from common off-target transduction of cells around the PBN, a line with broad expression (*Adcyap1*) was injected in the regions dorsal and rostral to the PBN (**Figure 8D-E**). This resulted in widespread expression in the CUN, ICe, ICc, and NLL, with only a few cells in the rostral PBls. The result was strong expression in the VMH, geniculate nucleus, and PAG (particularly the dorsal lateral PAG). There were also weak projections to the LS, MPO/LPO region, PVH, AD, CM, RE, LHA, much of the hypothalamic regions, DMH, medial amygdala, PF, ZI, VTA, RPA, MARN, RN, and NTS. Consequently, care must be taken to closely examine transduction of cells outside the PBN to avoid misinterpretation of results.

## DISCUSSION

The goals of this project were to establish a catalog of neuronal cell types within the PBN and characterize their transcriptional profiles, determine their location within the PBN, and map their axonal projections to facilitate studies of their functions in responding to and relaying interoceptive and exteroceptive signals. As expected, the round tissue punches used to isolate the PBN included some adjacent regions; consequently, 5 of the 21 clusters are not in the PBN. About 90% (6855/7635) of the PBN neurons are glutamatergic and most of them have discrete locations within the PBN, whereas the GABAergic neurons are scattered throughout. Some of the excitatory neurons (e.g., *Calca, Pdyn, Tacr1*) were already known to occupy distinct locations within the PBN and have distinct functions. Twelve of the PBN glutamatergic clusters have discrete locations. Our results reveal that most of the PBN neurons express multiple neuropeptides and GPCR receptors; the co-expressed transcription factors may play an important role in their cell-specific gene expression. Many of neuropeptides expressed in the PBN, e.g., cholecystokinin, substance P, calcitonin gene-related peptide, somatostatin, prodynorphin, neurotensin, where known to be expressed in the PBN based on immunohistochemistry studies and confirmed by *in situ* hybridization (Hermanson et al., 1998, Saleh and Cechetto, 1996, Shimada et al., 1985) The overall axonal projections of the PBN neurons were established using anterograde tracers and confirmed using S*lc17a6^Cre^*mice and injections of AAV carrying Cre-dependent fluorescent genes (Huang et al., 2021a). We provide detailed projection profiles from the PBN for many additional Cre-driver lines of mice that may generate ideas for testing their functions. The current data set provides a baseline for examining how the PBN changes during development and in response to environmental threats ranging from acute noxious events to chronic adverse conditions.

The PBN was classically divided into 10 sub-regions based on morphological criteria, anterograde and retrograde mapping strategies. While most studies agree with the relative location of these subregions their precise boundaries are not established; it is unfortunate that the names of the subregions and their abbreviations are inconsistent. We used the AMBA nomenclature because it is readily available, and frequently used. Gene expression patterns can help refine the boundaries of these sub-regions as shown in **Figure 4**. We identified genes that are primarily restricted to each of the PBN subregions except the PBme. Despite the tight clustering of some mRNAs, e.g., *Calca*, in specific sub-regions of the PBN, nearly all of them are also expressed more sparsely in neighboring regions. Furthermore, what was once considered to be a unique population turns out to be more complex, e.g., both the *Calca* and *Pdyn* neurons are represented by two clusters based on scRNA-Seq analysis. Whether these closely related clusters have distinct projections and functions needs to be established, e.g., are distinct *Pdyn* neurons involved in conveying temperature, nocifensive behaviors and feeding responses? The significance of the tight clustering of *Calca* neurons, with few if any interspersed neurons, is not known; it is unlikely that these neurons are connected by gap junctions to allow simultaneous activation because *Calca* neurons are not activated synchronously when GCaMP signals of individual neurons were analyzed (Chen et al., 2018).

Most PBN glutamatergic neurons are primarily restricted to a subregion with a few neurons scattered in other regions. In contrast, GABAergic neurons are rare (∼10% of total) and they are dispersed throughout the PBN, in agreement with 12% inhibitory neurons in rat PBN (Raver et al., 2020). Although GABAergic neurons in the KF project axons to the brainstem (Geerling et al., 2017), it is not known whether other PBN neurons project outside the PBN. Chemogenetic or optogenetic activation of GABAergic neurons in the PBN can inhibit the function of local *Slc17a6* neurons (Sun et al., 2020) and we have shown using electrophysiology techniques that their activation can inhibit *Calca^tdT^* neurons (unpublished).

There are several limitations to the conclusions reached in this study. The choice of parameters used for clustering of scRNA-Seq data can affect the number of clusters and sequencing depth can influence the reliability of co-expression for low-abundance transcripts. For example, the number of cells in a neuronal subcluster expressing a particular GPCR mRNA (e.g., *Oprm1*) was low based on scRNA-Seq analysis, whereas the signal from the Cre-driver line revealed many cells. We obtained ∼100,000 reads per neuron which is close to the number of mRNA molecules/cell. A higher number of reads is necessary to capture rare transcripts since a single transcript can maintain ∼10,000 proteins with a half-life of 1 day, which may be enough for many regulatory proteins. There was good, low-resolution correspondence among HiPlex, AMBA, and fluorescent protein expression from Cre-driver lines when transcripts were abundant, but *in situ* failed for some probes. When the distinguishing transcripts were of low abundance/cell, we were unable to detect any neuronal signal by *in situ* hybridization (e.g., *Fn1, Mylk, Slc35d3, Shisal2b*); this could be due to poor quality probes or expression levels that were too low to detect. In some cases, the probes were good but there was little signal in the PBN (e.g., *Nfib, Pappa*). There were also cases where predictions of transcript co-expression were not supported by HiPlex analysis. Cre-driver lines are superior for locating some neurons with low levels of gene expression (e.g., *Ntsr1, Chat, Avpr1a*) because, in principle, the action of a pair of Cre recombinase molecules is sufficient to activate robust expression from AAV carrying a Cre-dependent gene. When genes are expressed in the PBN as well as neighboring regions, restricting AAV transduction to the PBN is challenging. Interpretation of the axonal projections requires analysis of multiple injected mice and knowledge of where neighboring neurons project their axons.

The last decade has seen extensive use of Cre-driver lines of mice and AAV carrying Cre-dependent effector genes to interrogate the functions of PBN neurons. One strategy uses Cre-drivers with restricted expression. For example, numerous studies with *Calca^Cre^, Pdyn^Cre^, and Tacr1^Cre^* mice have revealed their activation by a wide variety of real and potential threats, while their optogenetic or chemogenetic activation is generally aversive and their inactivation ameliorates aversive responses to threats (references are included in Results, under **N10, N15, N17**). An alternative strategy uses Cre-driver mice with widespread expression in the PBN, e.g., *Slc17a6^Cre^, Cck^Cre^, Oprm1^Cre^,* and *Tac1^Cre^*, to assess behavioral/physiological consequences of their activation or inhibition (Barik et al., 2018, Cheng et al., 2020, Chiang et al., 2020, Liu et al., 2022, Sun et al., 2020). A problem with these Cre-drivers is the difficulty in restricting viral transduction to the PBN. Both approaches have flaws because it is rare for threats to activate only one cell type and all the cells represented by the more widely expressed genes are rarely engaged by specific threats. Furthermore, activation of some PBN neurons can suppress the activity of others, e.g., activation of *Tac1* neurons counteracts the outcomes of activating *Calca* neurons even though *Tac1* and *Calca* are extensively co-expressed (Arthurs and Bertsch, to be submitted). This possibility complicates interpretation of results when groups of neurons are artificially manipulated.

One solution to this problem is tagging and manipulating of groups of neurons that are normally engaged by specific threats, e.g., by using FosTrap, CANE or similar strategies (DeNardo and Luo, 2017, Sakurai et al., 2016), a technique that has been used to trap PBN neurons activated by pain and aversive odors (Liu et al., 2022, Rodriguez et al., 2017).

The broad outlines of axonal projections from the PBN were established decades ago by anterograde tracing studies (Fulwiler and Saper, 1984, Gauriau and Bernard, 2002, Krout and Loewy, 2000, Moga et al., 1990, Norgren and Leonard, 1971, Tokita et al., 2009) and confirmed by using broadly expressed *Slc17a6^Cre^* mice and injection of AAV expressing Cre-dependent fluorescent markers (Chiang et al., 2020, Huang et al., 2021a). The axonal tracts to the forebrain follow two main paths as they leave the PBN. The CTT pathway travels to the forebrain via the ventral thalamus, to the extended amygdala and cortex. The VP travels through the ventral tegmental area to the hypothalamus, with a periventricular branch passing through the periaqueductal grey and more dorsal thalamus. A descending pathway innervates parts of the hindbrain including the nucleus of the solitary tract (NTS), pre-Bötzinger and reticular regions. *Calca* neurons have axon collaterals that go to more than one brain region (Bowen et al., 2020). Most of the Cre-driver lines tested here resemble either the *Calca* neurons or the *Pdyn* neurons in their axonal projections; however, there are distinct differences in their innervation of forebrain targets that is revealed by synaptophysin staining. We identified two unique projections: one from *Phox2b* cells in the PBlc and PBls that innervate the SH and another from *Ntsr1* cells just outside the PBN in the NLL that innervate the VMH. As expected, reporter expression from Cre-driver lines with expression in several PBN clusters have composite projection patterns.

The axonal inputs to specific clusters of PBN neurons are beginning to be established, either by retrograde rabies virus tracing studies starting with specific Cre-drivers, e.g., *Calca^Cre^* (Liu et al., 2022, Rodriguez et al., 2017) or by candidate approaches starting with expression AAV-DIO-ChR2 into distal sites of Cre-driver lines of mice and recording photoactivated currents in specific PBN neurons, e.g., input from *Cck* or *Dbh* neurons in the NTS to *Calca* neurons in the PBN (Roman et al., 2016), *Oxt* neurons in pre-optic area to *Oxtr* neurons in PBN (Ryan et al., 2017), *Slc17a6* or *Slc32a1* inputs from BNST to *Pdyn* and *Calca* neurons in PBN (Luskin et al., 2021). Many other molecularly defined inputs to different subregions of the lateral PBN have been described, e.g., *Tacr1* and *Gpr83* from spinal cord (Choi et al., 2020), *Gfral, Glp1r* from area postrema (Zhang et al., 2021), *Calcr* from NTS (Cheng et al., 2020), *Slc17a6/Dbh* from LC (Yang et al., 2021), *Slc6a3* from ventral tegmental area (Han et al., 2021), *Slc32a1* from substantia nigra reticulata, *Npy/Slc32a1* from arcuate nucleus (Alhadeff et al., 2018, Wu et al., 2009); *Mc4r* from the PVN (Garfield et al., 2015), *Htr2a, Prkcd,* or *Sst/Pdyn/Crh* from CEA (Cai et al., 2014, Douglass et al., 2017, Raver et al., 2020). In each of these cases, knowing the locations and identity of PBN clusters that they innervate would refine connectivity maps and provide greater insight into potential functions.

Our study highlights the diversity of cell types in the PBN, many of which reside in distinct subregions that align with prior anatomical tracing studies. We also mapped the PBN cell distribution and detailed brain-wide projections of 21 Cre-driver mouse lines. All these data are publicly available for download and most of the mice have already been deposited at The Jackson Laboratory. We hope that this rich resource will continue to inspire future studies on the role and neurocircuitry of the PBN.

## Supporting information

Video 1 - Calca

## MATERIALS AND METHODS

### Key resources table

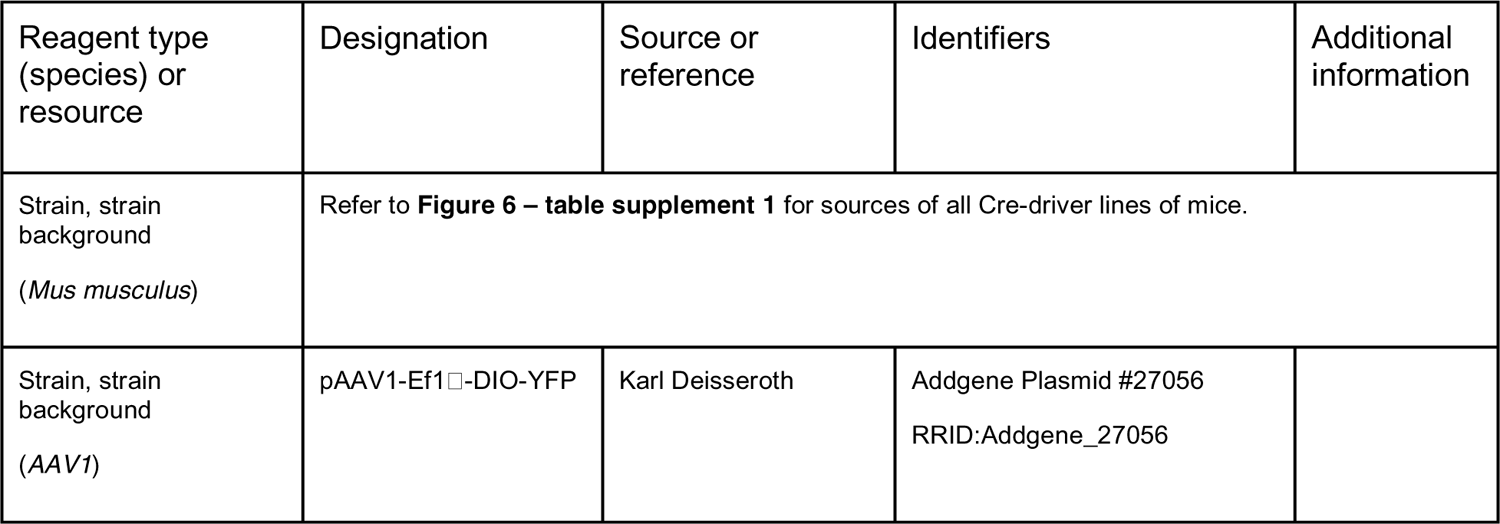
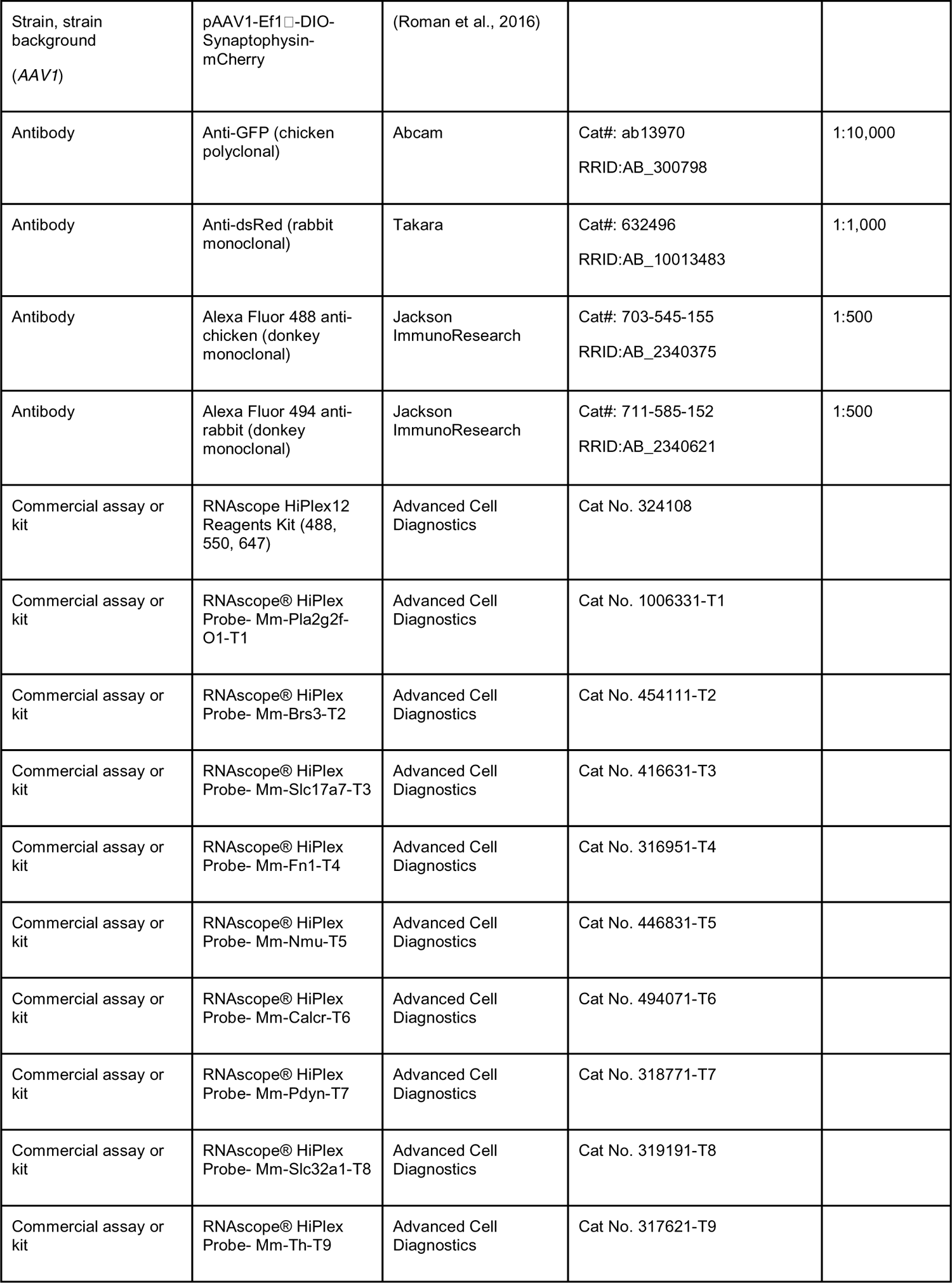
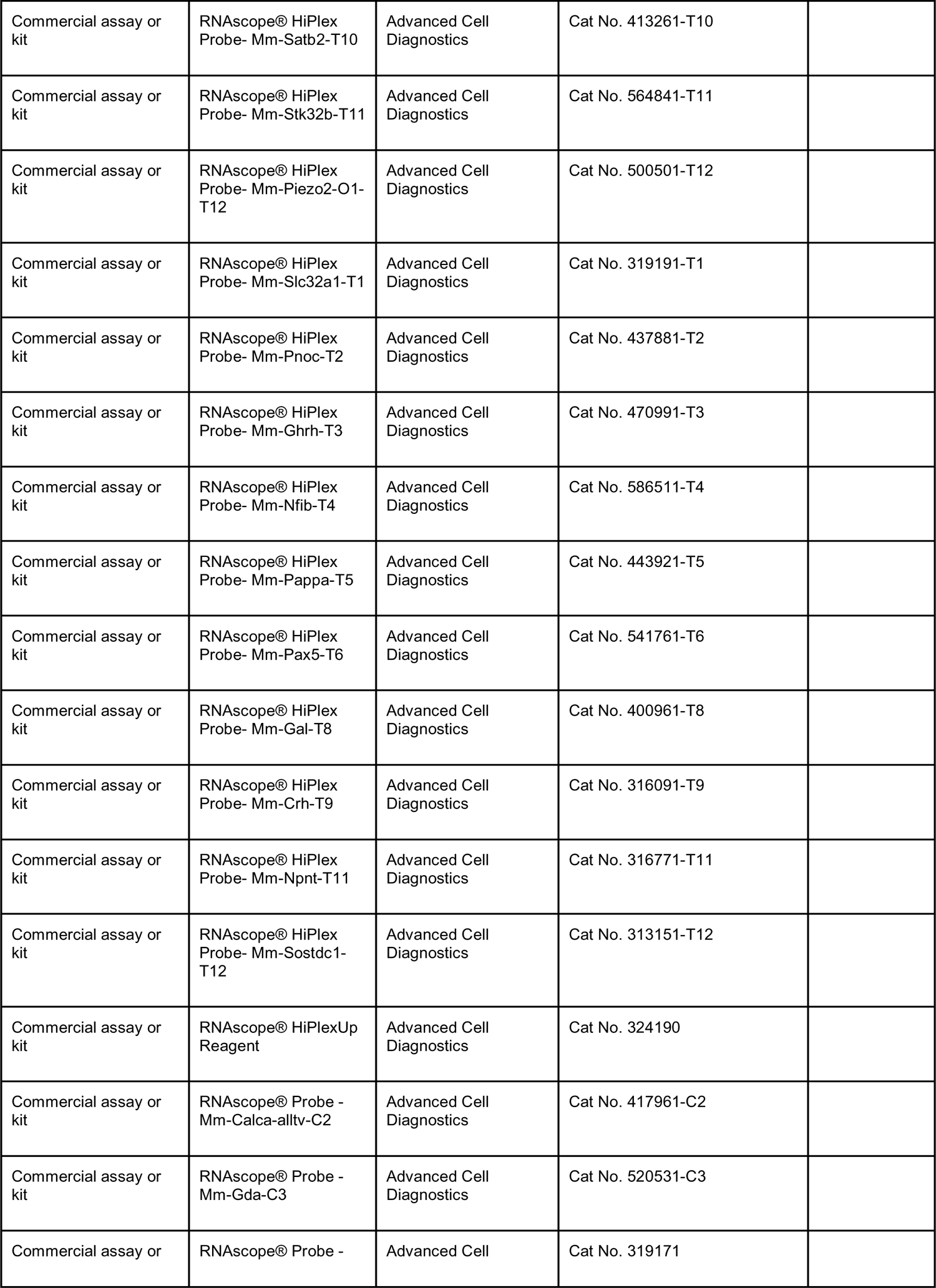
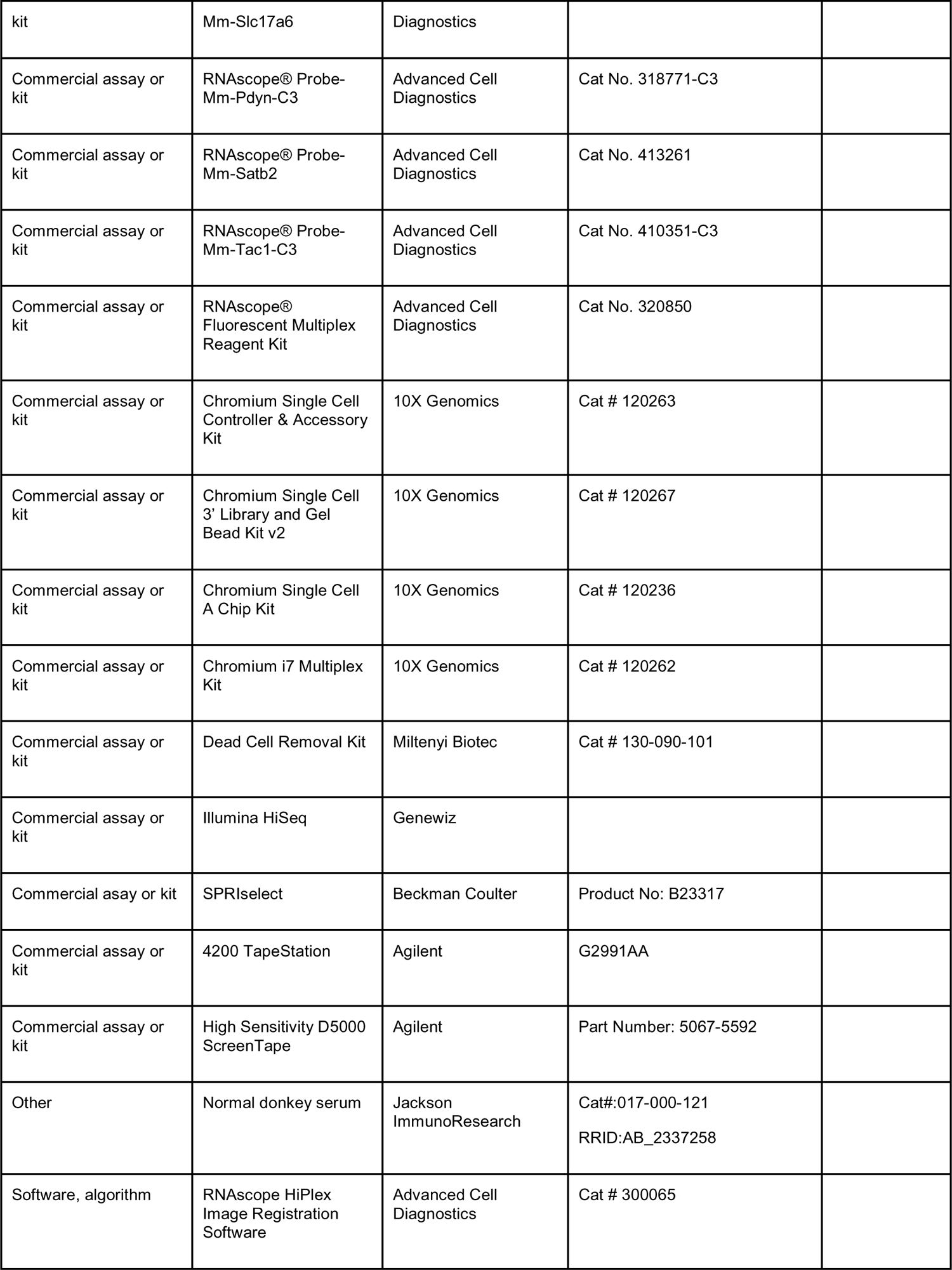
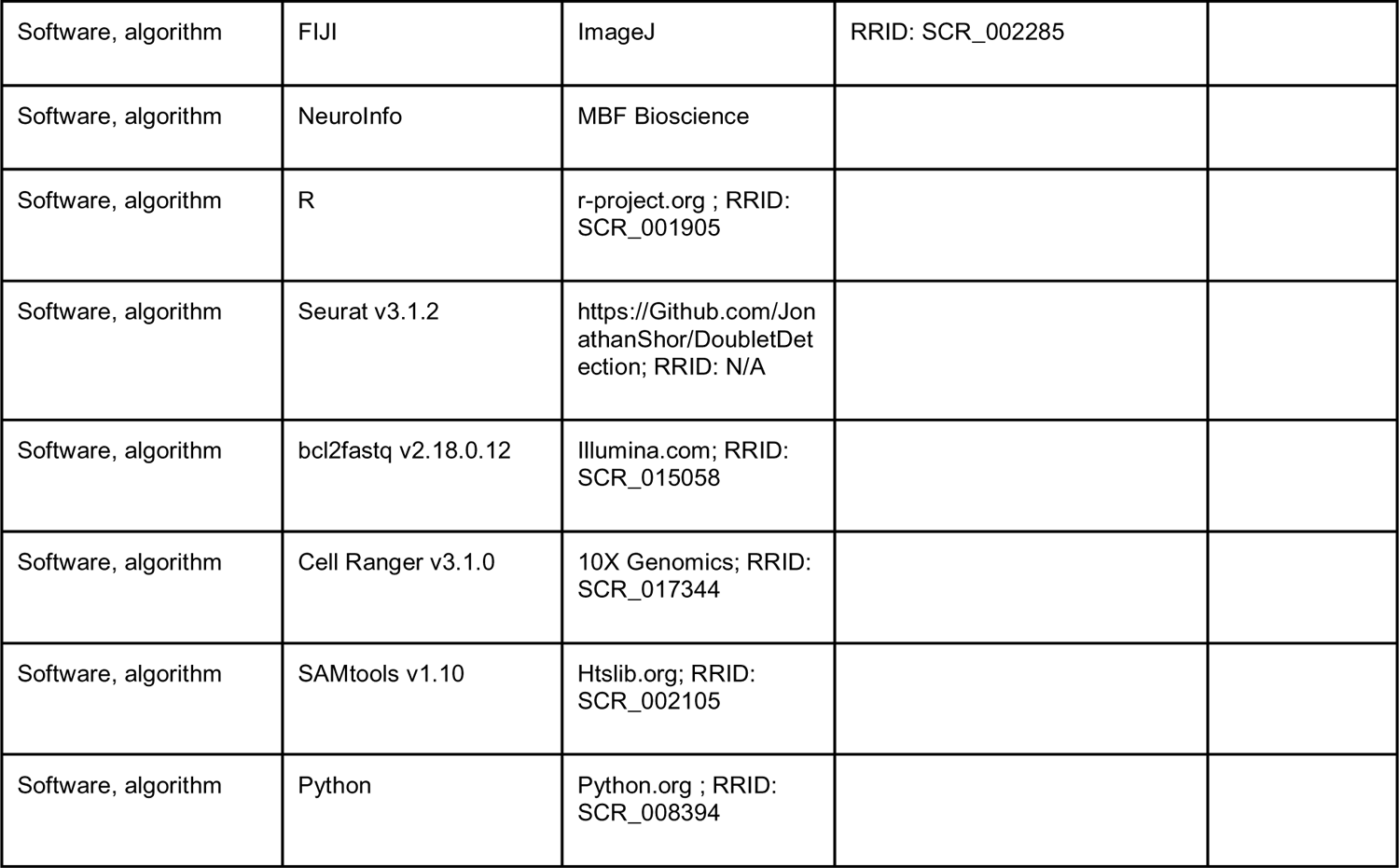

#### Animals

All experiments were approved by the Institutional Animals Care and Use Committee at the University of Washington. Animals were group-housed with littermates on a 12-h light cycle at ∼22°C with food and water available *ad libitum*. Male and female mice from the same litter were used, but no formal comparisons were done between sex.

#### Single-Cell Library Preparation and Sequencing

Live single cell suspensions were prepared as previously described (Rossi et al., 2021, Rossi et al., 2019) and tissue was harvested approximately 6 h after the onset of the dark cycle (Zeitgeber time ∼18:00). Briefly, mice were transcardially perfused with a cold artificial CSF solution containing NMDG (NMDG-aCSF), modified from (Ting et al., 2018). All steps were performed under continuous oxygenation and CO_2_ buffering using 95%/5% O_2_/CO_2_. Brains were rapidly dissected, and coronal slices (200 µm) spanning the PBN were prepared using a vibrating microtome (VT1200, Leica Biosystems). Slices were allowed to recover in NMDG-aCSF containing 500 nM TTX, 10 µM AP-V, 10 µM DNQX (NMDG-aCSF-R) for 20 min and the PBN was subsequently isolated using tissue punches (500-750 μm). The isolated tissue was enzymatically digested using 1 mg/mL Pronase (Roche) for 50 min, and enzymatic digestion was quenched with 0.05% BSA. Cells were mechanically dissociated using a fire-polished glass pipet with an internal diameter of 200-300 µm, filtered through a 40 μm strainer, washed, and depleted of dead cellular fragments using a commercial kit (Miltenyi Biotec, Bergisch Gladbach, Germany). The remaining cells were resuspended in PBS containing 0.05% BSA at a concentration of 1,000 cells/µL.

Single-cell RNA libraries were generated using Chromium Single-Cell 3’ v2 chemistry (10X Genomics, Pleasanton, CA) following the standard manufacturer protocol. A pool of ∼17,000 cells harvested from 5 mice were loaded per reaction, with the first pool run on a single reaction and the second pool spread across three reactions **(Figure 1 – figure supplement 1H-I)**.

For each library, cDNA amplification was performed using 12 cycles and indexing was performed using 11 cycles. Library size distribution and concentration was determined using a Qubit HS DNA Assay (Invitrogen, Waltham, MA) or High Sensitivity D5000 ScreenTape (Agilent Technologies, Santa Clara, CA). Each reaction library was sequenced on two lanes of an Illumina HiSeq 4000 using 2×150 chemistry by Genewiz, Inc (South Plainfield, NJ) following the standard 10X Genomics v2 paired-end configuration. Approximately 875 million reads were generated per lane with a mean sequencing saturation of 82.6%. Sequences were aligned to the mm10-3.0.0 genome, and digital expression matrices were created using 10X Genomics Cell Ranger v3.1.0 with 128 GB of memory on 24 cores.

### Single-Cell Clustering Analysis and Feature Discovery

Clustering was performed using Seurat v3.1.2 (Stuart et al., 2019) and custom code in R v3.6.1 as described (Rossi et al., 2021). Briefly, to remove low-quality cells, total input cells were first filtered using a threshold for genes and fraction mitochondrial reads, and doublets were subsequently removed using DoubletDetection v2.5.4 under default parameters on Python 3.7 via Reticulate v1.14 (Gayoso and Shor, 2022). Cells containing ≤ 800 genes and ≥ 10 percent mitochondrial reads were removed from the primary analysis (**Figure 1** and **Figure 1 – figure supplement 1)**. Within the analysis of all cells, one subcluster was unable to be mapped to any specific features and was excluded from analysis (**Figure 1B, gray)**. Following these filters, a total of 39,649 cells were retained with a median of 1,740 genes and 3,366 transcripts represented across a median of 47,177 reads per cell (**Figure 1** and **Figure 1 – figure supplement 1A-E)**.

The neuronal cluster identified in the initial analysis of all cells (**Figure 1B**) was then isolated and subjected to a more stringent quality threshold where only cells containing ≤ 2 percent mitochondrial reads were retained to enable high-confidence, high-resolution sub-clustering **(Figure 2 – figure supplement 2C** and **H).** Within the neurons, two subclusters were unable to be mapped to any specific features and were excluded from the analysis (**Figure 2A, gray)**. This resulted in a final analysis of 7,635 neurons containing a median of 3,189 genes, 7,823 transcripts, and 99,583 reads per cell (**Figure 2** and **Figure 2 – figure supplement 1A-J)**.

Regression and integration of samples was performed using a regularized negative binomial regression (Hafemeister and Satija, 2019, Stuart et al., 2019) and canonical correlation analysis (Butler et al., 2018, Stuart et al., 2019) as described (Rossi et al., 2021) **(Figure 1 – figure supplement 1F-G** and **Figure 2 – figure supplement 1K-M)**. Principal components were calculated on all genes, and cells were plotted in UMAP space using iteratively tuned parameters to optimize for visualization (McInnes et al., 2018) (**Figure 1 – figure supplement 1J** and **Figure 2 – figure supplement 1O**). Clustering was performed using the Louvain algorithm with multilevel refinement (Rodriguez and Laio, 2014) with a K parameter of 25 and resolution of 0.04 (all cells) (**Figure 1**), or a K parameter of 15 and resolution of 0.25 (neurons) (**Figure 2**). Feature discovery for cell-type assignment was performed on Pearson residuals using a likelihood-ratio test for single-cell data as implemented in Seurat (Hafemeister and Satija, 2019, Macosko et al., 2015, McDavid et al., 2013).

Prepossessing, regression, integration, dimensionality reduction, clustering, and feature discovery were run on a Dell blade-based cluster at the University of North Carolina at Chapel Hill running Linux RedHat Enterprise 7.7. All other steps were performed on an Apple MacBook Pro running macOS 11.4.0.

Raw and processed data for the scRNA-seq experiment has been deposited at the National Center for Biotechnology Information Gene Expression Omnibus (NCBI GEO, accession number GSE207708; https://www.ncbi.nlm.nih.gov/geo/query/acc.cgi?acc=GSE207708). The code used in this analysis is available at a Github repository affiliated with the Stuber Laboratory group (https://github.com/stuberlab/Pauli-Chen-Basiri-et-al-2022).

### RiboTag Analysis of Transcripts Enriched in *Calca* Neurons

The *Calca^Cre^* mice were used to identify mRNAs that were selectively being translated in neurons that express the *Calca* gene by injecting a virus expressing a Cre-dependent hemagglutinin (HA)-tagged ribosomal protein 22 (AAV-DIO-Rpl22-HA) (Sanz et al., 2015) into the PBN of 8 adult mice. After several weeks to allow incorporation of the tagged ribosomal protein into ribosomes, tissue punches (two pools of bilateral punches from 4 mice) were collected, total cell extract was prepared and then polyribosomes were precipitated with an antibody against the HA-tagged ribosomes (Sanz et al., 2019, Sanz et al., 2009). For microarray analysis, 10 ng of total RNA was amplified and biotin-labeled using the Ovation Pico SL WTA system with the EncoreIL biotinilation module (NuGEN), and 750 ng of the labeled cDNA was hybridized to a MouseRef-8v2.0 gene expression BeadChip (Illumina). Signal was detected using the BeadArray Reader (Illumina) and analyzed using the GenomeStudio software (Illumina). Average normalization and the Illumina custom error model were applied to the analysis. Only transcripts with a differential score of >13 (p < 0.05) were considered.

Raw and normalized RiboTag data have been deposited in the NCBI GEO (accession number GSE207153; https://www.ncbi.nlm.nih.gov/geo/query/acc.cgi?acc=GSE207153). In some cases, the microarray included more than one probe for the same gene and the absolute values could differ greatly; thus, the relative enrichment is more reliable.

### Mice

The *Calca-tdTomato* mice have been described (Jarvie et al., 2021). All the Cre-driver lines of mice used in this study are listed in **Figure 6 – table supplement 1**. Most of them have been described previously and/or deposited at Jackson labs, where details can be found. All mice were on a C57BL/6J genetic background. The generation of 4 new lines are described below. The *Cbln4, Ntsr1,* and *Tacr1* Cre-driver lines were made by inserting IRES-Cre:GFP just beyond the termination codon of these genes, whereas the *Pdyn* line was made by inserting Cre:GFP just 5’ of the initiation codon. The 5’ and 3’ arms (5 to 8 kb) were amplified from a C57Bl/6 BAC clone by PCR using Q5 polymerase (New England Biolabs) and inserted into a cloning vector that contains frt-flanked SV-Neo for positive selection and *Pgk*-DTA and HSV-TK genes for negative selection (Jarvie et al., 2021). The linearized construct was electroporated in G4 ES cells (129 x BL6). About 80 G418-resistant clones were picked and expanded for Southern blot analysis using a 32P-labeled probe located just beyond either the 5’ or 3’ arm. Correctly targeted clones were injected into blastocysts and transferred to recipient female mice. Germline transmission of the targeted allele was determined by 3-primer PCR (2 primers flanking the termination codon region and 1 reverse primer in the IRES). A single insert was confirmed by Southern blot. Mice were bred with *Rosa26-FLPe* mice to remove the SV-Neo gene and then bred with C57BL/6 mice for at least 6 generations.

### Stereotaxic Surgery and Axon-Projection Tracing

Mice were anesthetized with isoflurane and placed on a robotic stereotaxic frame (Neurostar GmbH, Tübingen, Germany) and AAV-1EF1a-DIO-YFP and AAV1-EF1a-DIOsynaptophysin:mCherry were injected bilaterally into the PBN (AP −4.8 mm, ML +/- 1.4 mm, DV 3.5 mm) at a rate of 0.1 µl/min for 2 min. At least 3 weeks after virus injection, mice were deeply anesthetized with sodium pentobarbital and phenytoin sodium (0.2 ml, i.p.) and intracardially perfused with ice-cold PBS followed by 4% PFA. Brains were post-fixed overnight in 4% PFA at 4 °C, cryoprotected in 30% sucrose, frozen in OCT compound, and stored at −80 °C. Coronal sections (35 µm) spanning the brain (Bregma 2.0 mm to −8.0 mm) were cut on a cryostat and collected in cryoprotectant for long-term storage at −20°C.

Sections were washed two times in PBS and incubated in a blocking solution (3% normal donkey serum and 0.2% Triton X-100 in PBS) for 1 h at room temperature. Sections were incubated overnight at 4°C in blocking solution with primary antibodies including: chicken-anti-GFP (1:10000), and rabbit-anti-dsRed (1:2000). After 3 washes in PBS, sections were incubated for 1 h in PBS with secondary antibodies: Alexa Fluor 488 donkey anti-chicken and Alexa Fluor 594 donkey anti-rabbit, (1:500). Tissue was washed 3 times in PBS, mounted onto glass slides, and coverslipped with Fluoromount-G with DAPI (Southern Biotech).

Whole-slide fluorescent images were acquired using a Keyence BZ-X710 microscope and higher magnification images using an Olympus FV-1200 confocal microscope and minimally processed using Fiji to enhance brightness and contrast for optimal representation of the data. For TIFF stacks, images were aligned using the BrainMaker workflow in NeuroInfo (MBF Bioscience).

### RNAscope Multiplex/HiPlex FISH

Mice were deeply anesthetized with sodium pentobarbital and phenytoin sodium (0.2 ml, i.p.), decapitated, and brains rapidly frozen on crushed dry ice. Coronal sections (15 µm) were cut on a cryostat, mounted onto SuperFrost Plus slides, and stored at −80°C. RNAscope HiPlex Assay or RNAscope Fluorescent Multiplex Assay were performed following the manufacturer’s protocol.

Images centered on the scp in the PBN were acquired in a 3×3 grid at 20x using a Keyence BZ-X710 microscope and stitched together 4 sets of in 4 channel stacks using Fiji. Images of the probe staining within the 4-channel sets were subtracted from one another using Fiji’s image calculator function to remove background autofluorescence. The Dapi images from each of the 4 sets of images were registered using the HiPlex Image Registration Software (ACDBio) and then used to register all the probe images. This process was repeated for a second HiPlex experiment. Colors were assigned using the registration software.

### Evaluation of Gene Expression and Projection Density

Registered HiPlex probe images were combined into 5 stacks for each bregma level for both experiments. PBN and surrounding subregions of interest (ROIs) were drawn based on the AMBA designations and distinct probe locations. Using these ROIs, each probe was scored based on an estimation of the number of transcripts present per cell and number of cells labeled per region. Projection regions were evaluated based on density by matching the brain sections in the TIFF stacks as closely to the AMBA as possible.

## Data Availability

Raw and preprocessed data for scRNA-seq: NCBI GEO accession number GSE207708; https://www.ncbi.nlm.nih.gov/geo/query/acc.cgi?acc=GSE207708

Code for analysis of scRNA-Seq data: https://github.com/stuberlab/Pauli-Chen-Basiri-et-al-2022

Raw and normalized data for RiboTag: NCBI GEO accession number GSE207153; https://www.ncbi.nlm.nih.gov/geo/query/acc.cgi?acc=GSE207153

Images from RNAscope and all tracing experiments: Zenodo DOI: 10.5281/zenodo.6707404; https://doi.org/10.5281/zenodo.6707404

## Author Contributions

Conceptualization: RDP

Investigation: JLP, JYC, MLB, SP, MEC, ES

Data Curation: JLP, JYC, MLB

Writing – Original Draft: RDP

Writing – Review & Editing: JLP, JYC, MLB, SP, ES, GDS, RDP

Visualization: JLP, JYC, MLB

Supervision: RDP, GDS, GSM

Project Administration: JYC, RDP

Funding Acquisition: RDP, GDS, GSM

## Acknowledgments

We thank Susan Phelps for maintaining the mouse lines used in these studies. This work was supported in part by grants from the National Institutes of Health, R01- DA24908 (R.D.P), R01-DA032750 and R01-DA038168 (G.D.S).

---

**Figure 1 – figure supplement 1.**
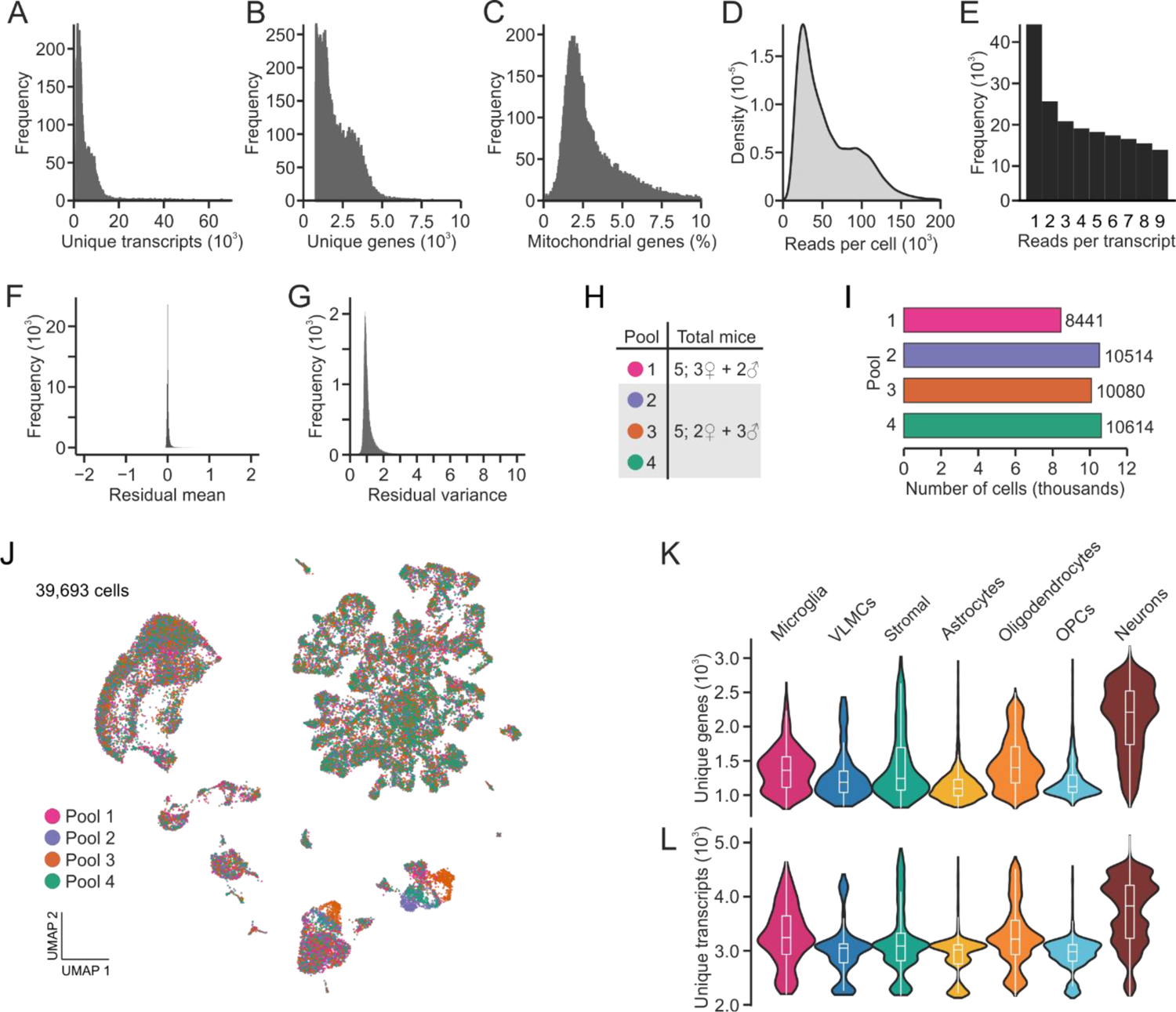
Technical metrics in scRNA sequencing analysis of resident PBN cell types. **(A)** Distribution of unique transcripts per cell. **(B)** Distribution of unique genes per cell. **(C)** Distribution of percent mitochondrial reads per cell. **(D)** Distribution of total sequencing reads per cell. **(E)** Distribution of reads per transcript. **(F)** Following integration, the mean of residuals centers on zero. **(G)** Following integration, mean variance centers on one. **(H)** Number and sex of mice used in each library pool. **(I)** Number of cells sequenced from each library pool. **(J)** Following integration, each pool is represented uniformly across UMAP space. **(K)** Distribution of unique genes across each cell type **(L)** Distribution of unique transcripts across each cell type.

**Figure 2 – figure supplement 1.**
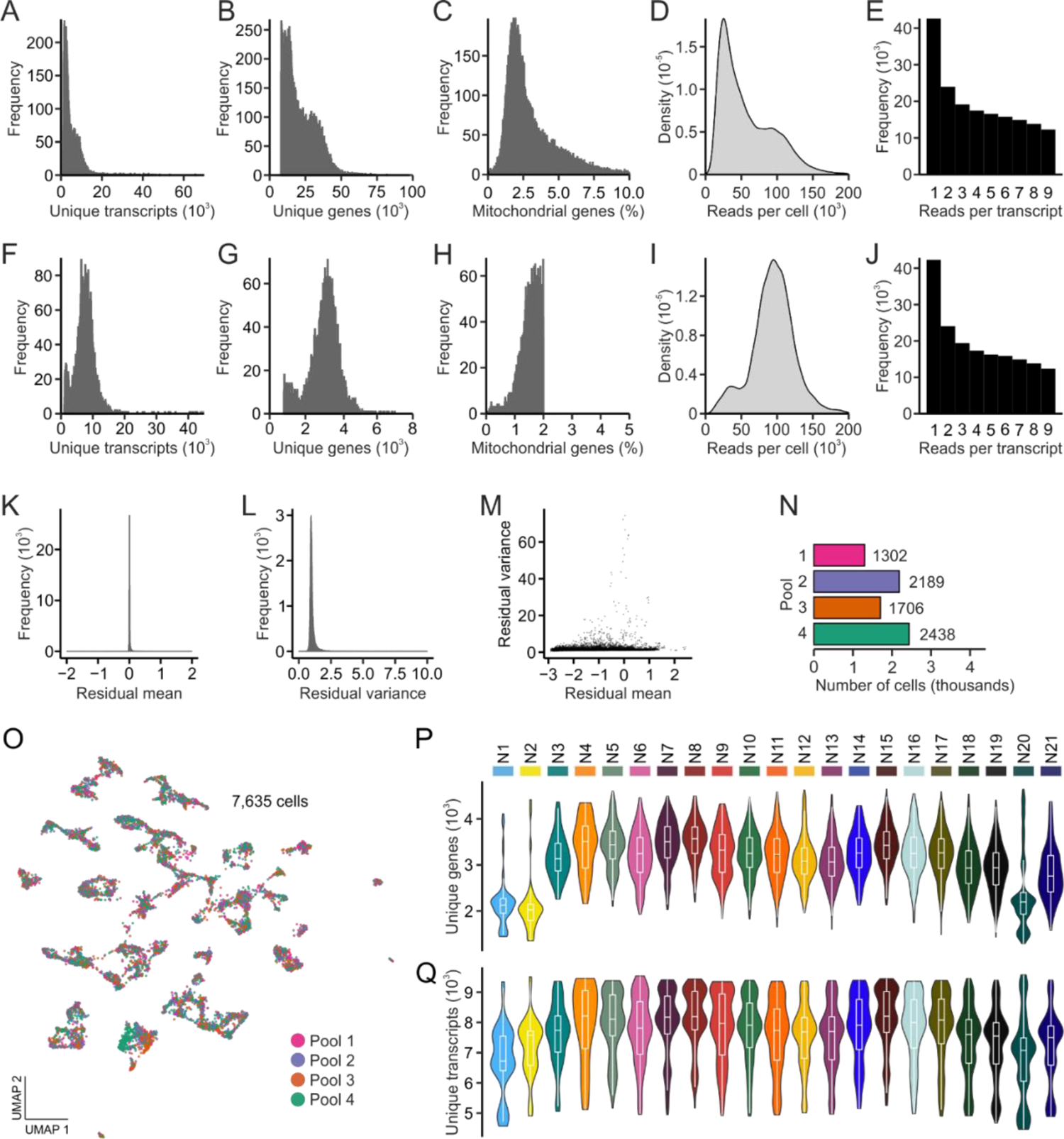
Technical metrics in scRNA sequencing analysis of neuronal subclusters. **(A)** Distribution of unique transcripts per cell. **(B)** Distribution of unique genes per cell. **(C)** Distribution of percent mitochondrial reads per cell. **(D)** Distribution of total sequencing reads per cell. **(E)** Distribution of reads per transcript. **(F)** Distribution of unique transcripts per cell after thresholding. **(G)** Distribution of unique genes per cell after thresholding. **(H)** Distribution of percent mitochondrial reads per cell after thresholding. **(I)** Distribution of total sequencing reads per cell after thresholding. **(J)** Distribution of reads per transcript after thresholding. **(K)** Following integration, the mean of residuals centers on zero. **(L)** Following integration, mean variance centers on one. **(M)** High-variance residuals were assessed for clustering analysis. **(N)** Number and sex of mice used in each library pool. **(O)** Following integration, each pool is represented uniformly across UMAP space. **(P)** Distribution of unique genes across each neuronal subcluster **(Q)** Distribution of unique transcripts across each neuronal subcluster.

**Figure 3 – table supplement 1.**
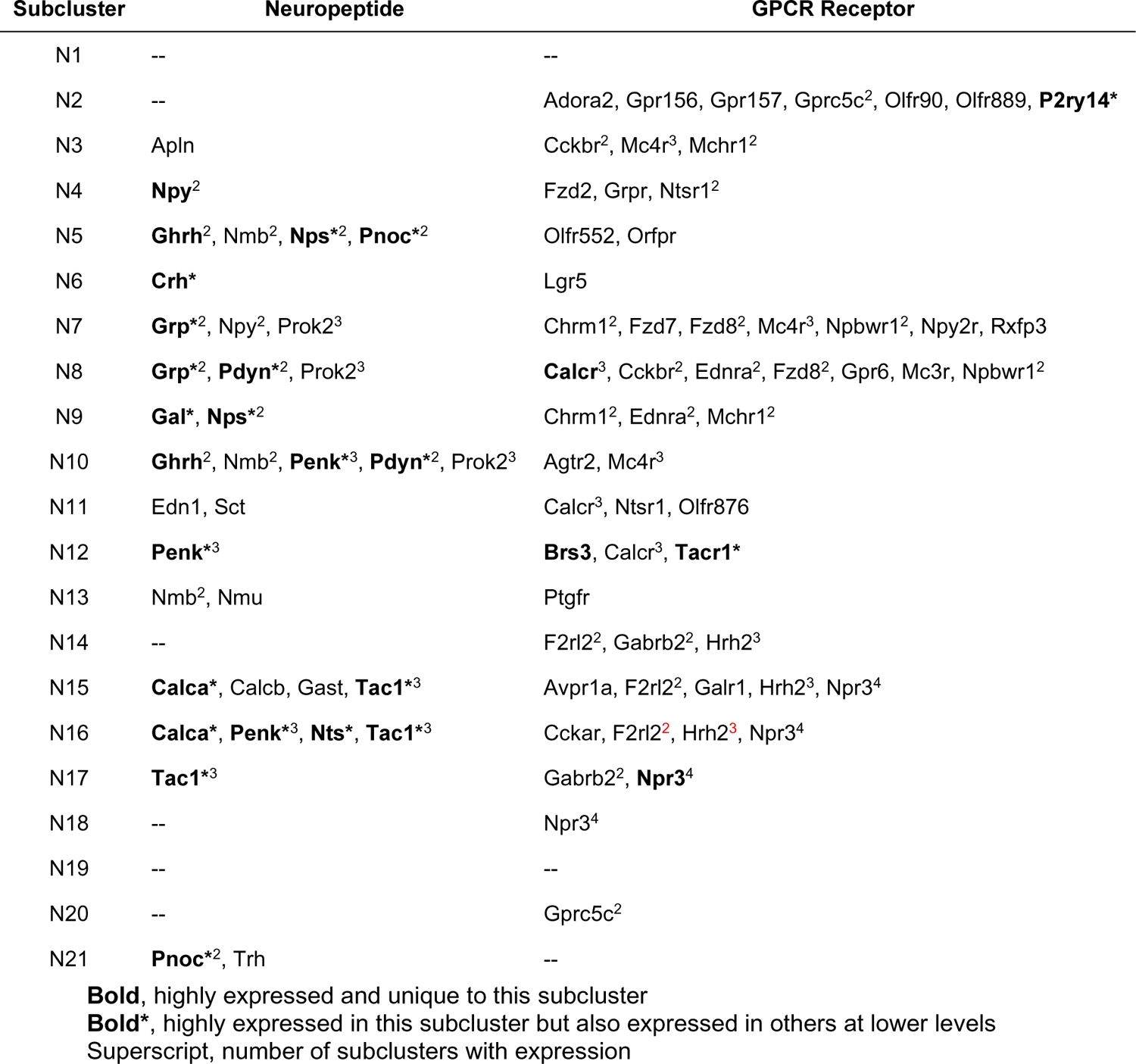
Neuropeptides and GPCRs with restricted expression in the 21 neuronal clusters. These data are extracted from **Supplementary File 3** and **Supplementary File 4**.

**Figure 4 – figure supplement 1.**
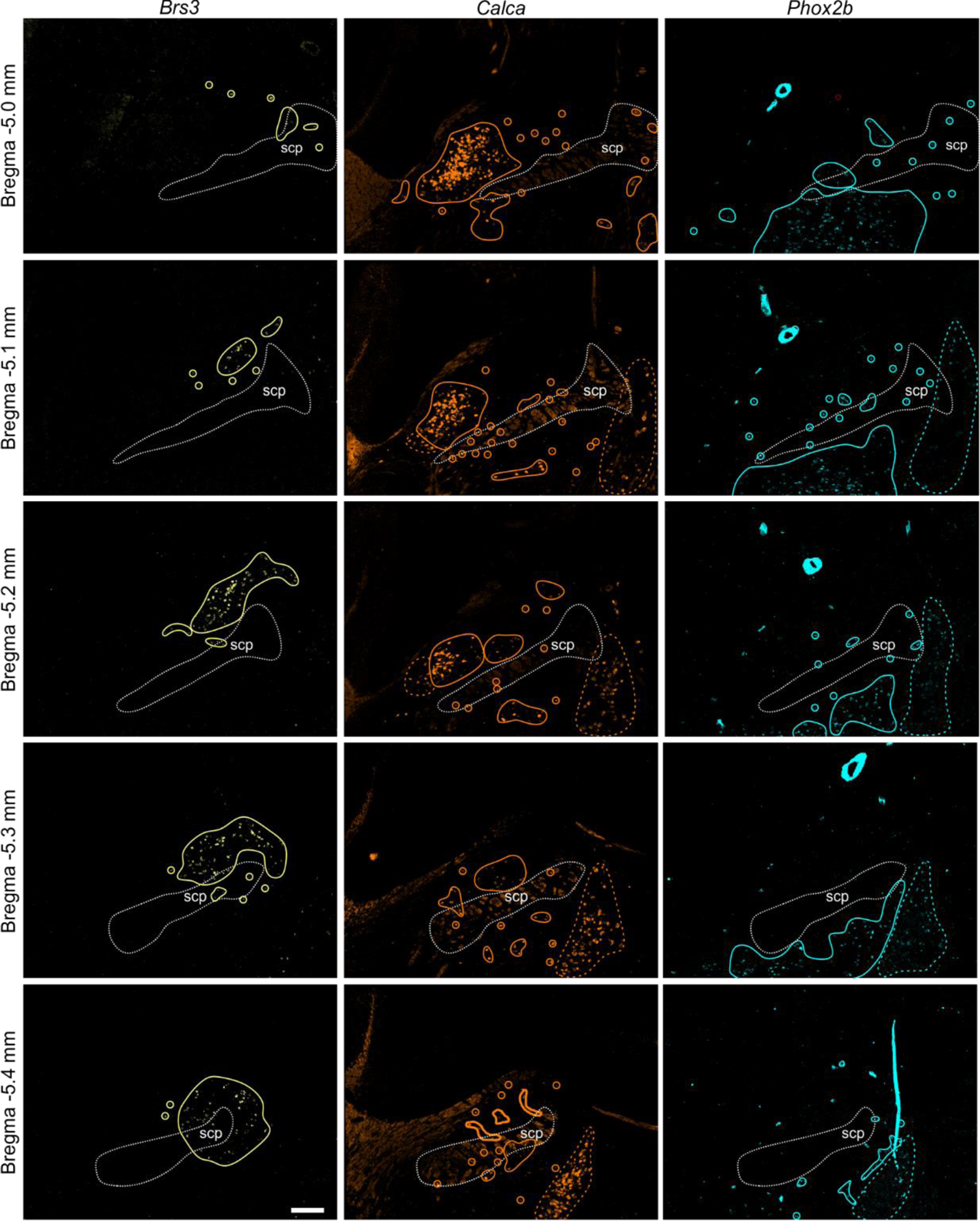
Example of HiPlex staining for *Brs3*, *Calca*, and *Phox2b* for 5 Bregma levels. Solid lines surround clusters of positive neurons or individual neurons, colored dashed lines indicate expression outside the PBN, such as in KF and LC. Scale bar, 200 μm.

**Figure 5 – figure supplement 1a.**
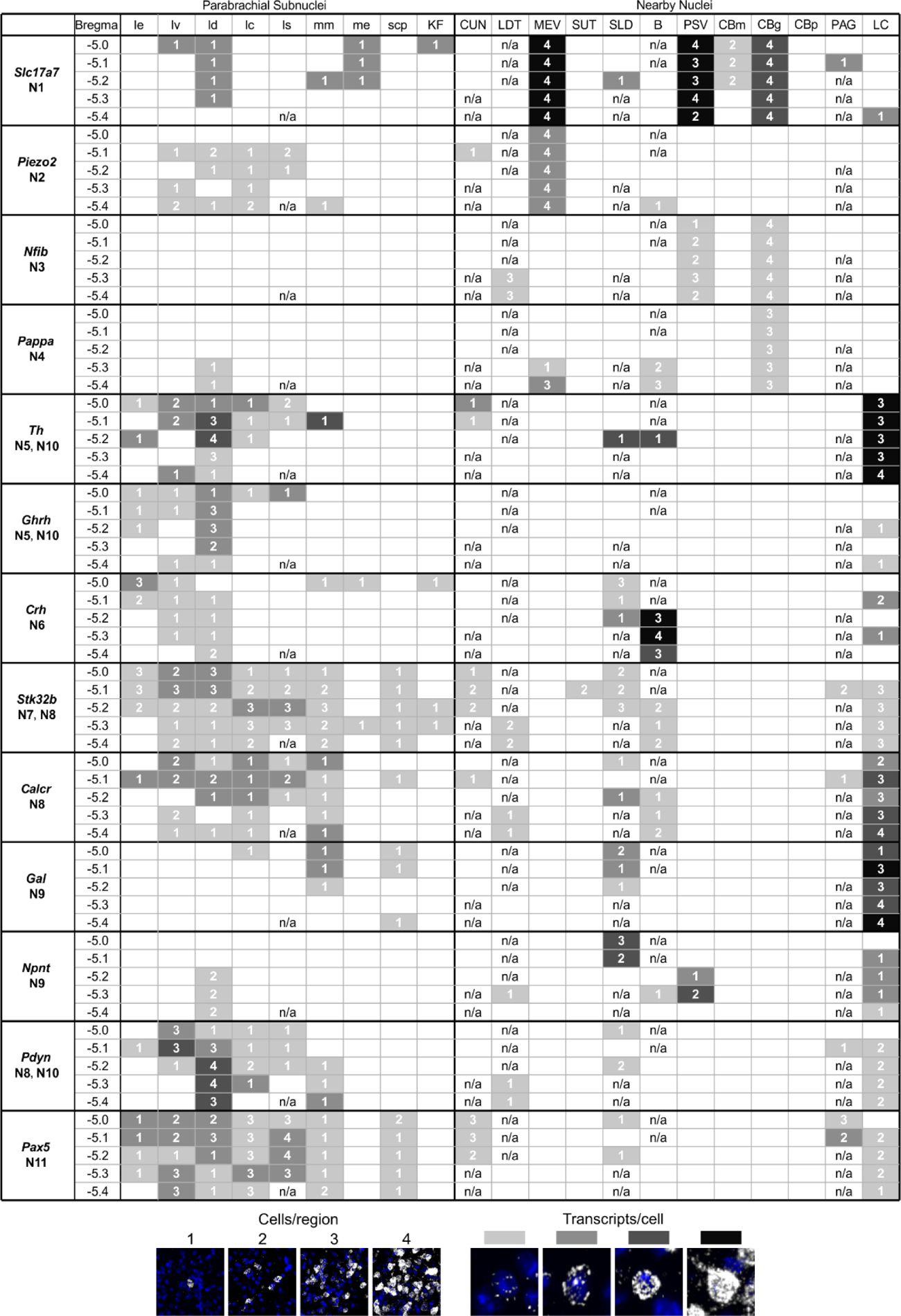
HiPlex results for all probes with signal in PBN and surrounding area, page 1.

**Figure 5 – figure supplement 1b.**
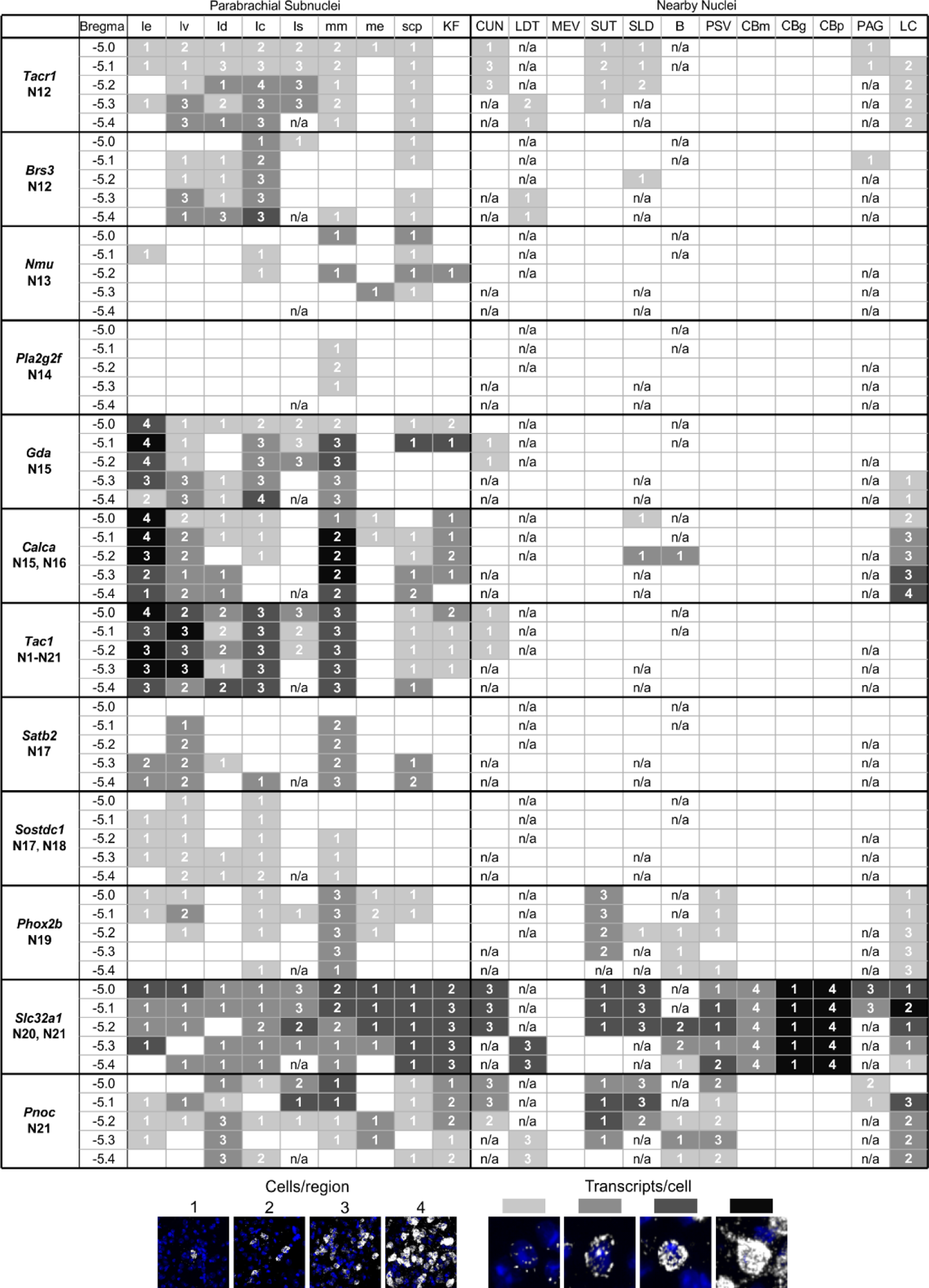
HiPlex results for all probes with signal in PBN and surrounding area, page 2.

**Figure 5 – figure supplement 2.**
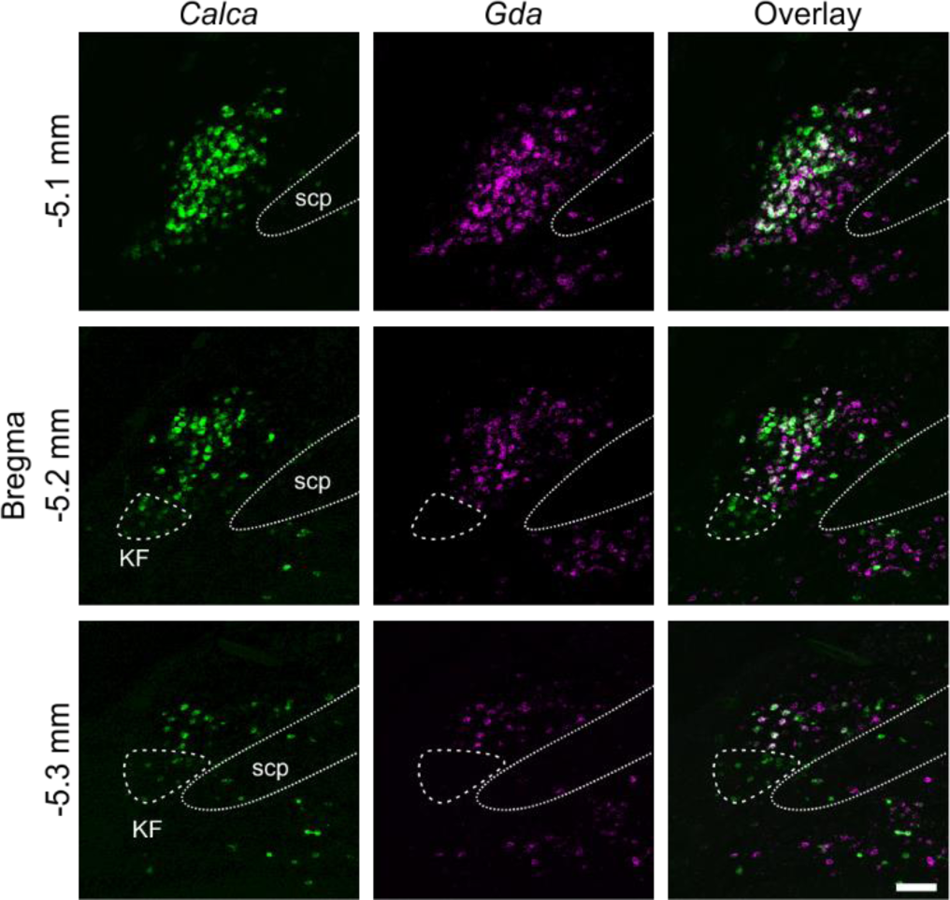
RNAscope for *Calca* and *Gda*. Co-expression represents **N15** based on scRNA-Seq data. *Calca*-only cells that do not express Gda (circled by dashed line) are located in the ventral lateral PBle that extends partially into the Koelliker-Fuse (KF) represent **N16**. Scale bar, 100 μm.

**Figure 5 – figure supplement 3.**
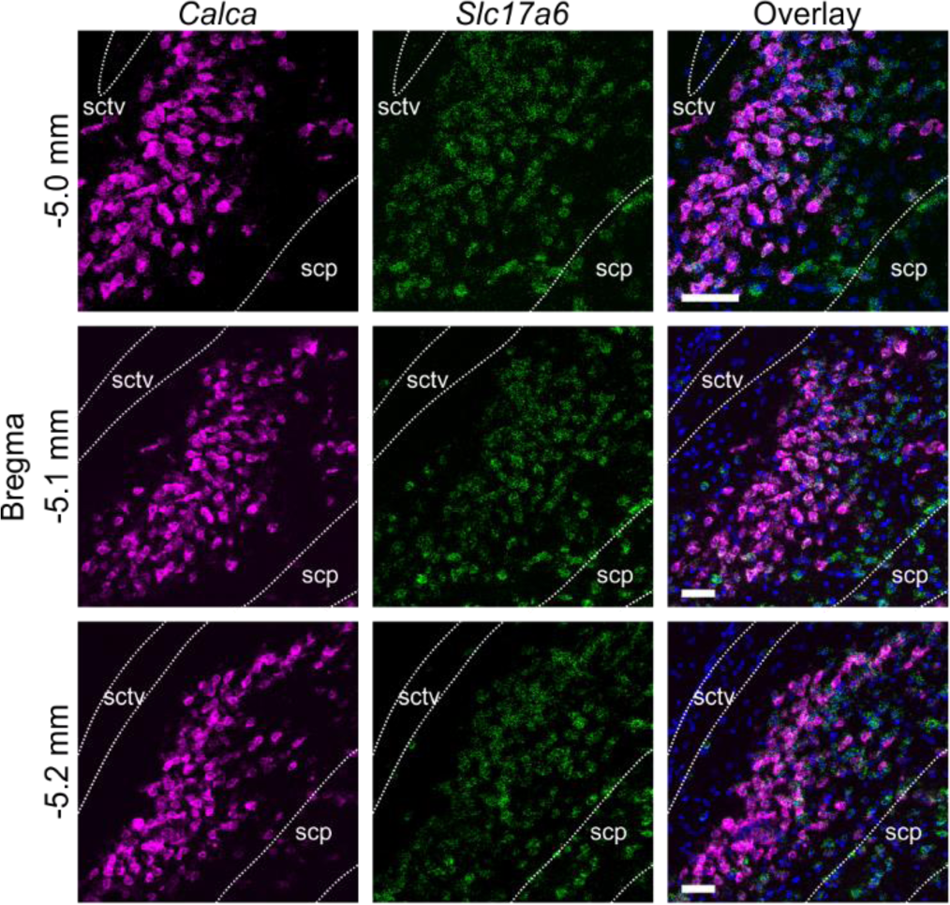
RNAscope for *Calca* and *Slc17a6*. Nearly all *Slc17a6*-positive cells in the core of PBle also express *Calca*. Dapi is shown in blue in the Overlay image. Scale bar, 100 μm.

**Figure 5 – figure supplement 4.**
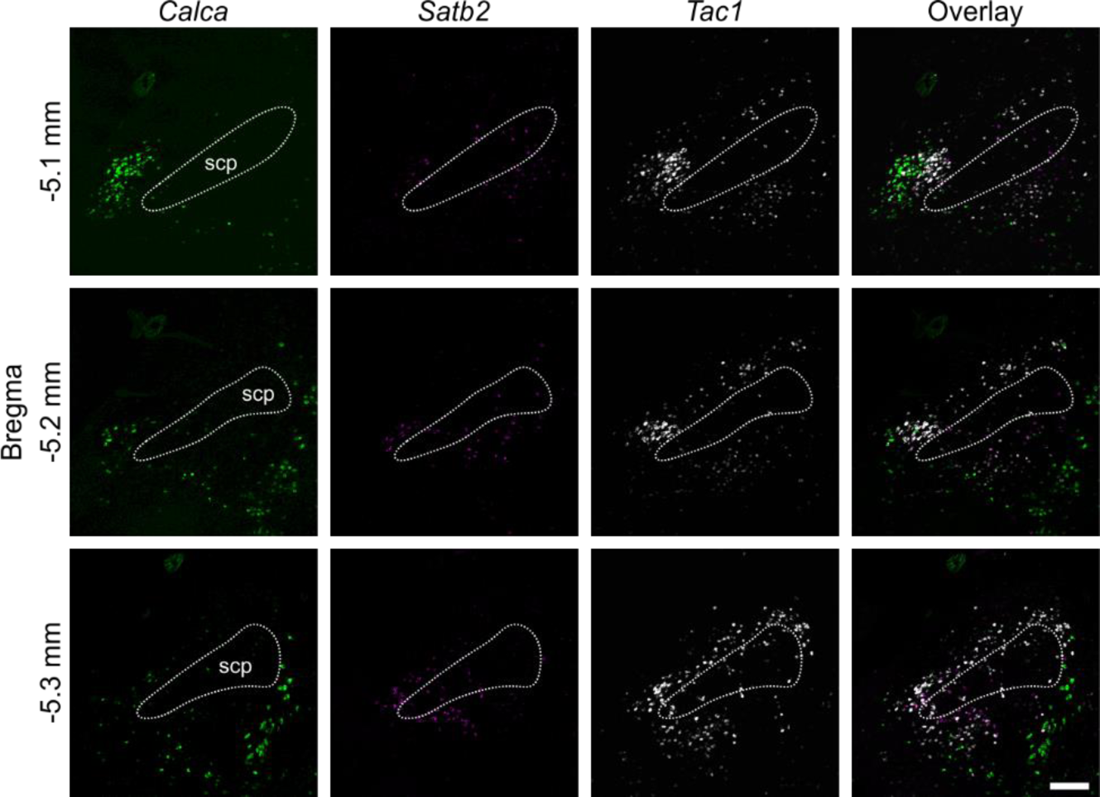
RNAscope for *Calca*, *Satb2* and *Tac1*. Difference in expression patterns of *Calca*, *Satb2* and *Tac1*. Scale bar, 200 μm.

**Figure 5 – table supplement 5.**
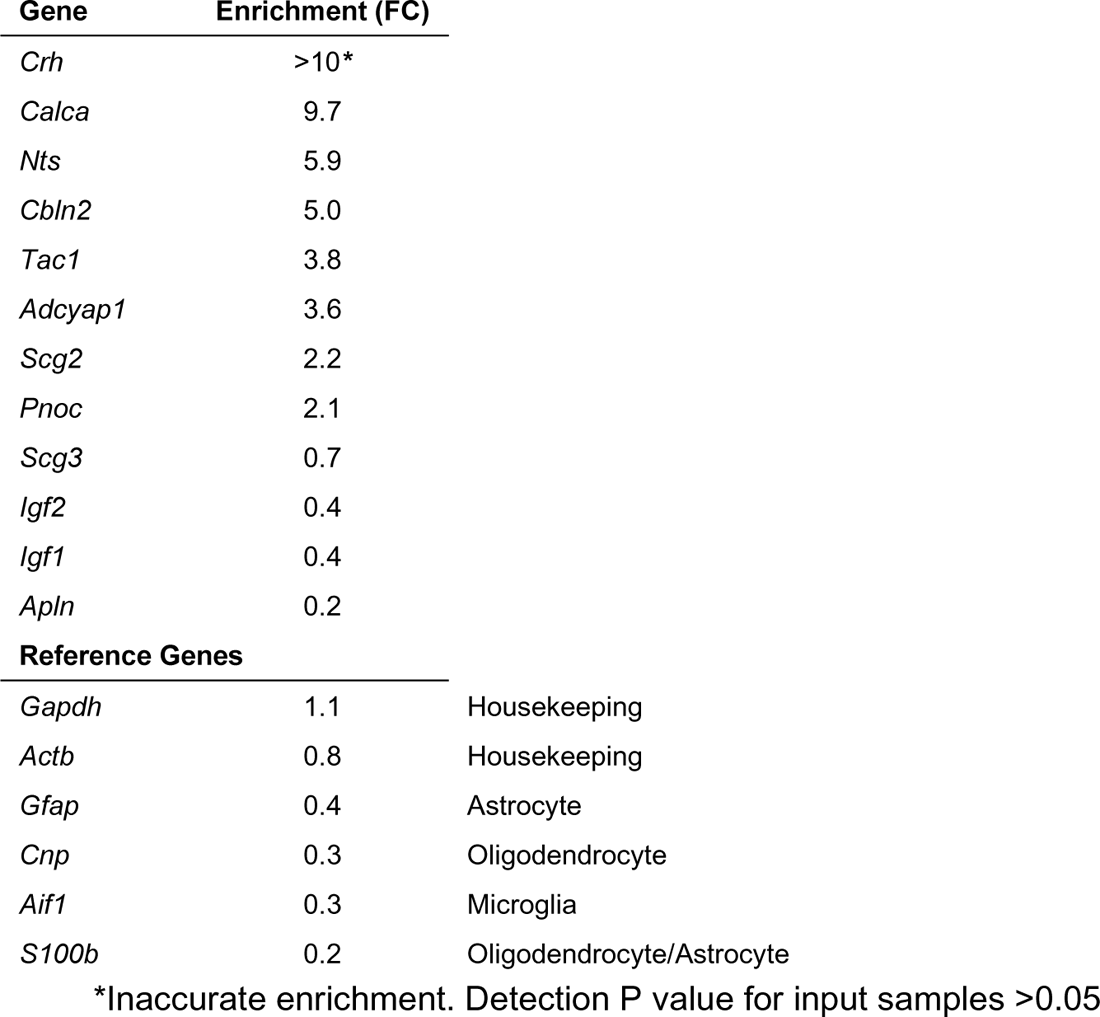
Enrichment of neuropeptide mRNAs in *Calca* neurons based on RiboTag experiment, measured by the ratio of immunoprecipitate to input (FC, fold change). Only significantly different (p<0.05) neuropeptide mRNAs are shown. Housekeeping and glial mRNAs are included for reference. **Supplementary File 6.** Source data for Ribotag experiment showing all genes (1) and genes significantly enriched/depleted (p<0.05) sorted by fold change (FC, 2).

**Figure 6 – table supplement 1.**
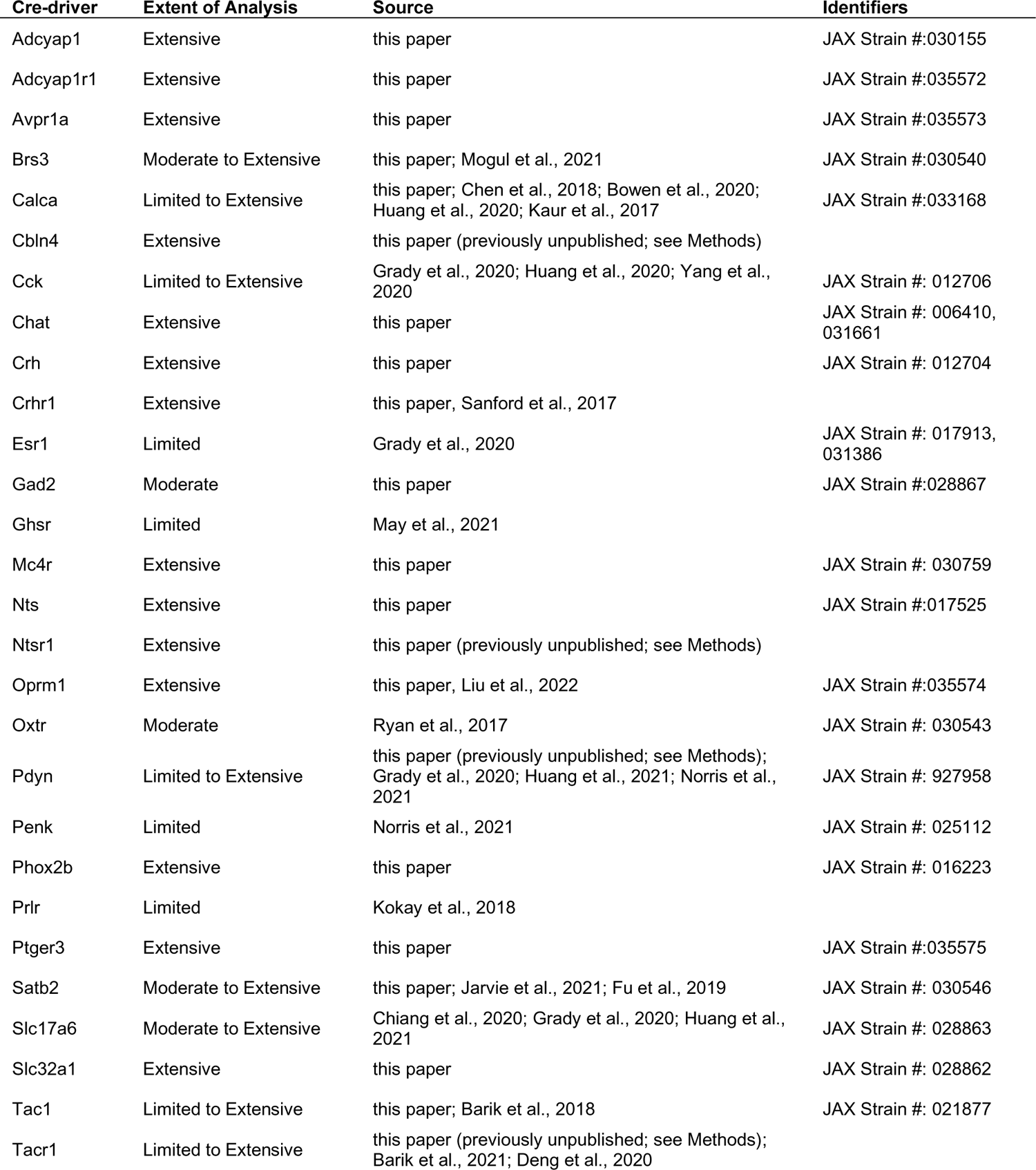
Table of all Cre-driver lines of mice that have been used to study expression in PBN and axonal projections.

